# On the conditions for shifts in metabolic strategies

**DOI:** 10.1101/2025.10.03.680276

**Authors:** Maarten J. Droste, Robert Planqué, Frank J. Bruggeman

## Abstract

Many heterotrophic microorganisms gradually replace an energetically-efficient mode of metabolism by an inefficient, more-wasteful overflow metabolism above a critical growth rate, even though the energy demand continues to rise with growth rate. For instance, complete respiration of a sugar is replaced by its fermentation. We derive the conditions for this to happen, using a core model of metabolism and growth that is qualitatively in agreement with experimental data. Assuming a fixed cellular protein content, the model shows that protein expression of efficient metabolism and anabolism rises as function of growth rate until a critical value is reached. This growth-associated protein expression is at the expense of proteins associated with future adaptation. At the critical growth rate, this preparatory pool is reduced to zero. Beyond the critical growth rate, the anabolic protein pool and the energy demand continue to rise and therefore less protein remains for catabolism. In this regime, the inefficient metabolism gradually takes over ATP synthesis from the efficient mode. It can do so if it requires less protein per unit of ATP flux. The catabolic protein pool then decreases while the anabolic pool increases. This continues until all catabolic protein is allocated to the inefficient strategy and to anabolism. We show that this can occur only if the maximal growth rate of the inefficient mode is higher than the critical growth rate. We show that this condition is precisely the requirement for the second mode to be more proteome-efficient than the first mode. Finally, we reduce a genome-scale model of protein expression in the yeast *Saccharomyces cerevisiae* to a variant of our core model and show that it is still qualitatively in agreement with the experimental data used to validate the original model.

## 1 Introduction

The conceptual framework of studying microbial physiology from the perspective of limited biosynthetic resources for protein expression aims to understand protein expression behaviour of (microbial) cells as function of conditions [1; 2; 3]. It was pioneered by several works, seminal ones are Maaløe and Kjeldgaard [4], Scott et al. [5] and Molenaar et al. [6]. A key aspect of this framework is that the increase of protein expression of one cellular task is invariably at the expense of another [7]. This principle applies when the performance of cellular tasks increases with protein investments, and single proteins are involved in unique tasks. Both apply generally, since proteins are often catalytic enzymes whose activity is proportional to their concentration, and involvement in multiple tasks is indeed rare.

A corollary of this principle is that key cellular functions trade off against each other, such as growth, alternative-nutrient adaptation, and stress tolerance [7; 8]. Indeed, fast-growing cells have been shown to be less adaptive and stress-tolerant than slow-growing cells [9; 10]. This happens because faster growth requires more growth-associated proteins since the needed higher metabolic rates are (predominantly) achieved by higher protein concentrations [11]. Since cellular macromolecules consist of the same building-block molecules and their polymerisation has a fixed energetic and biosynthetic cost [12], and since the carbon source requirements of alternative biosynthesis pathways of building blocks from precursor molecules in central carbon metabolism vary little [13], anabolic processes are largely constant across conditions; their protein investment is roughly proportional to their rates and therefore to the growth rate, across all conditions. Accordingly, anabolic protein pools indeed rise linearly with growth rate in experiments [14; 15].

Thus at faster growth more protein is allocated to anabolism, at the expense of protein for growth-unassociated processes (maintenance, new-nutrient adaptation and stress-tolerance) and for *catabolism*. Therefore, if alternative catabolic pathways exist that can each provide charged energy equivalents and precursor metabolites (for biosynthesis and polymerisation of building block metabolites) under the same conditions, then we could in principle expect a shift in the usage of catabolic pathways as function of growth rate.

One would expect that this shift occurs towards the (proteome-efficient) catabolic pathway that requires the least amount of protein for sustaining the required ATP synthesis rate for the current growth rate. (This ATP-synthesis with growth rate relation is given by the yield of biomass on ATP, set by the stoichiometry of the anabolic pathways, and can be expected to be constant.) The shift is expected to occur provided that the catabolic pathway outperforms the catabolic pathway that it replaces [16]. To understand such shifts, we therefore need to explain why this catabolic pathway is, in fact, not used throughout the entire growth rate regime. Moreover, we need to know which pathway properties allow us to predict whether a switch is expected [17]. Both are aims of this paper.

Shifts in catabolism as function of growth rate, in the manner just described, have been experimentally observed. The best understood and most referenced case is overflow metabolism, by, for instance, *Escherichia coli* [18; 19; 20] and other bacteria [21; 22; 23]. It also occurs in the yeast *Saccharomyces cerevisiae* [24; 25] and other Crabtree-positive yeasts [26; 27; 28]. This phenomenon occurs during aerobic growth in glucose-limited chemostats [25; 29], but also as function of the growth rate on different carbon sources in batch conditions [20]. In all these cases, respiration of a carbon source with carbon dioxide as catabolic product is traded in for a partial breakdown of the carbon source into catabolic products such as acetate, ethanol or lactate via overflow metabolism (e.g., fermentation). The second strategy harvests less ATP per unit carbon source, and is energetically more inefficient per unit of carbon; however, it is the preferred option at faster growth, even though more ATP is needed per unit time.

Such shifts in metabolism are not confined to gradual shifts from respiration to overflow metabolism: it can also happen that an efficient mode of fermentation is traded in for a different, less efficient, mode of fermentation that has a lower ATP yield per mole of the carbon source [30; 31]. An example of this is the shift of *Lactococcus lactis* from mixed-acid fermentation to homolactic fermentation that occurs in an aerobic, glucose-limited chemostat [32]. (This microbe lacks a respiratory chain.) Also acetogens display overflow metabolism [33]. Contrary to all previous examples, these microbes do not rely on glycolysis for ATP synthesis, but on the Wood-Ljungdahl pathway. Because of the widespread occurrence of overflow metabolism, we focus on a core model, phrased in general terms and the essential metabolic features.

The critical growth rate at which the shift from efficient to the inefficient mode of catabolism is initiated can be influenced by titrating the level of the growth-unassociated protein pool, as elegantly shown by Basan et al. [20]. Since this leaves the anabolic protein demand for a specific growth rate unaffected, it indicates that the shift occurs when insufficient biosynthetic resources remain for catabolism [20]. Our model therefore considers, in addition to kinetics and stoichiometry, the protein costs per unit flux of the associated metabolic pathways.

In this work, we provide a core model that allows for analytical solutions that are interpretable in simple terms. We show that the occurrence of overflow metabolism may be predicted by comparing the maximal growth rates of the different strategies the cell has available; the model applies both to chemostat cultures and to carbon uptake-limited growth in batch conditions, thus unifying our understanding. We also show how such a core model may be formulated by reducing a genome-scale model of metabolism used to predict protein concentrations. We apply this technique to model overflow metabolism in the yeast *Saccharomyces cerevisiae*. The resulting core model is shown to behave qualitatively in accordance to the same experimental data used for validation of the genome-scale model in Elsemman et al. [29].

## 2 Core model: definitions and optimisation

### 2.1 The core model

Metabolic shifts are studied by both large genome-scale metabolic models [16; 29; 34] and small phenomenological models [6; 20; 35; 36; 37]. Although large models contain more biochemical detail, both approaches yield (similar) explanations for the shifts [17]. The reason is that these models are all based on linear programming. The mathematical properties underlying metabolic shifts are thus preserved when condensing a large model to a small one. As long as the same number of (active) constraints and degrees of freedom are taken into account [38], the structure of the solution space of the model is not affected. The reason that a metabolic is observed is therefore retained, but may be studied with greater clarity in a smaller model.

The main ingredients for any of these models are (i) the stoichiometric matrix of the metabolic network; (ii) the assumption of steady state behavior of metabolism in balanced growth; (iii) the assumption that growth rate is maximised in any given condition; (iv) one or more limitations on cellular processes or resources. We now introduce a small core model featuring one pathway that makes efficient use of the carbon source, and one that is carbon-inefficient. Cells may use these pathways exclusively, or use both at the same time. A metabolic shift occurs if the cell changes between these different regimes.

The core model is depicted in Figure 1. It consists of four metabolic processes. Assimilation (*A*) converts the carbon and energy source S into the intermediate X^1^. The pathway *R* (e.g., respiration) converts X into the catabolic product P_*R*_ and produces *m*_*R*_ ATP per X. The pathway *F* (e.g., fermentation) converts X into the catabolic product P_*F*_ and produces *m*_*F*_ ATP per X. Finally, biomass is formed from X by consuming *m*_*B*_ ATP and *Y*_*X/B*_ X per unit biomass. Since protein synthesis is the major ATP consuming process during growth, we model biomass formation as protein synthesis. Accordingly, the sum of the cellular protein is denoted by *e*_*tot*_. The processes *A, R, F* and *B* represent lumped metabolic pathways.

**Figure 1.**
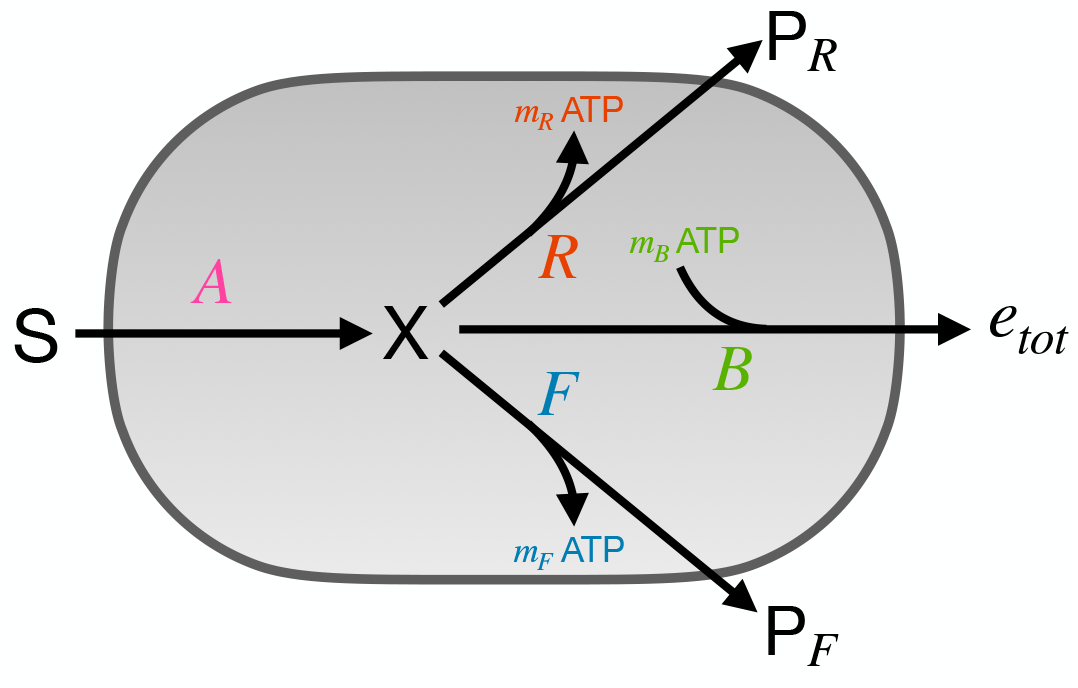
The coarse-grained metabolic network described by the core model. A carbon source S is assimilated (A) and converted into the intermediate X. Two catabolic modes (R, F) generate energy carriers (ATP) while converting X into catabolic waste products (P_*R*_, P_*F*_). Biomass synthesis (B) consumes the remaining carbon and generated ATP.

The core model depicted in Figure 1 is associated with the following differential equations for the concentrations of X and ATP,

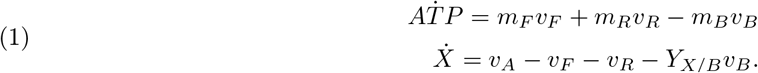

Denote the stoichiometric matrix by

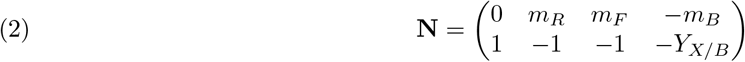

and the flux vector by

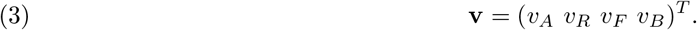

At balanced growth, metabolism is in steady state [39], i.e., **Nv** = **0**. The steady state flux vectors for this core model must have positive entries. The following two steady state vectors correspond to the modes using exclusively the *R* and the *F* pathways, respectively,

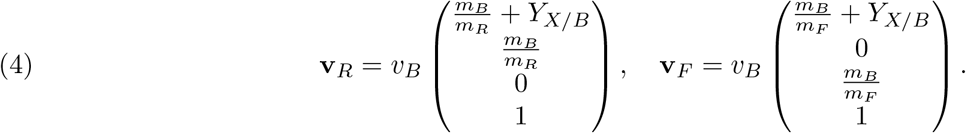

The *R* pathway is assumed to be most carbon-efficient, in the sense that it generates more ATP per unit of carbon: *m*_*R*_ *> m*_*F*_ . Later, we will also introduce the concept of proteome efficiency, which refers to the ATP synthesis rate per unit of protein. Both vectors have been normalised so that the biomass producing flux equals 1, and have the property that each reaction in the corresponding pathway is required to sustain a flux through it. Such flux vectors have been termed Elementary Flux Modes (EFMs) [40].

### 2.2 Optimisation of the growth rate

Explanations of overflow metabolism often invoke the assumption of growth rate maximisation. Single cells have an evolutionary advantage if their long term growth rate is high [8]. In constant conditions, this is equal to their steady state growth rate *λ*. The growth rate is bounded by limitations on cell growth: cells have limited biosynthetic resources, the total enzyme concentration in the cell is generally constant across conditions [15; 41; 42], and cells need to spend some of their enzymes on maintenance and preparation for future conditions such as changes in food availability or possible stresses [1; 8; 11].

To focus on these limitations, we introduce enzyme concentrations into the model, with each metabolic process represented by a reaction catalysed by its unique enzyme. Let *e*_*tot*_ be the total enzyme concentration, and assume it is constant across conditions. Then the rate of protein synthesis by ribosomes *v*_*B*_ balances with the rate of dilution by growth *λe*_*tot*_ [5],

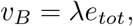

or in other words,

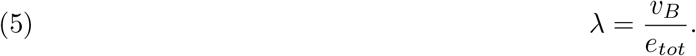

Each enzymatic reaction *j* has a corresponding enzyme concentration *e*_*j*_, catalytic rate constant *k*_*j*_ and saturation constant *f*_*j*_, with *v*_*j*_ = *k*_*j*_*e*_*j*_*f*_*j*_, e.g., *v*_*A*_ = *k*_*A*_*e*_*A*_*f*_*A*_. (In Supplementary Section S8 we show how one can estimate mean constants for a set of individual enzymatic reactions that are lumped into one process in a core model.)

In terms of protein concentrations, maximisation of the growth rate is formulated as an optimisation problem by

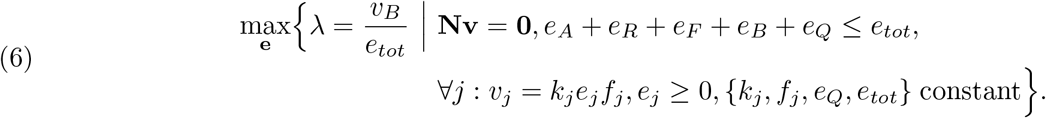

We have introduced one new protein pool, *e*_*Q*_, for maintenance processes, and assume it to be fixed across conditions [5]. Moreover, even though *e*_*tot*_ is assumed constant, the sum of the protein concentrations *e*_*A*_ + *e*_*R*_ + *e*_*F*_ + *e*_*B*_ + *e*_*Q*_ is not: this will be clarified in the next section.

The optimisation problem (6) can be rewritten in terms of the growth-associated fluxes *v*_*j*_ as variables

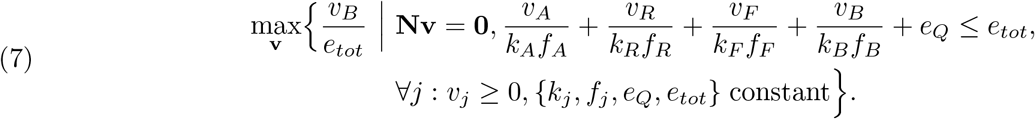

The set of admissible vectors may be characterised using the special EFM vectors **v**_*R*_ and **v**_*F*_ we defined in (4). Any positive steady state vector satisfies

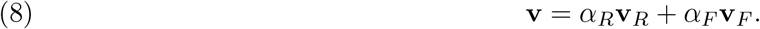

Here, *α*_*R*_ and *α*_*F*_ are convex coefficients that satisfy 0 ≤ *α*_*R*_, *α*_*F*_ ≤ 1 and *α*_*R*_ + *α*_*F*_ = 1.

In more general terms (i.e., for arbitrary stoichiometric matrices **N**), the set {**v** | **Nv** = **0, v** ≥ **0**} forms a pointed polyhedral cone that is spanned by its extreme rays [43]. These are called Elementary Flux Modes (EFMs) in the context of metabolic networks [40]. Therefore, any steady-state flux vector may be written as a convex combination of these rays. The EFMs are uniquely determined by the stoichiometric matrix **N**.

De Groot et al. [38] showed for this general case that the number of flux-carrying EFMs is bounded by the number of active flux-limiting constraints when a (specific) flux is optimised. We therefore infer (an upper bound on) the number of active EFMs by identifying the active constraints in the model. We thus know the root cause of the onset of overflow in general terms: if one adds extra constraints to (7), the maximiser switches from a single EFM to a mix of two or more. The downside of this formulation in terms of fluxes is that it is hard to interpret these additional constraints biologically; as a result, there are many putative explanations of overflow metabolism in the literature [17].

### 2.3 The core model for carbon-limited conditions

To provide a biological interpretation of the constraints in terms of actual limitations on cell growth, we adhere to the formulation in terms of protein concentrations (eq. (6)). The extra constraint follows from modeling different experimental conditions, such as carbon limitation. Various ways exist to realise this condition experimentally. During carbon uptake-limited batch cultivation, the carbon source is available in excess, but the experimentalist controls the carbon uptake flux by varying the nutrient quality and/or by titrating the expression of protein involved in nutrient assimilation (e.g. [20; 44]). The corresponding optimisation problem is formulated as

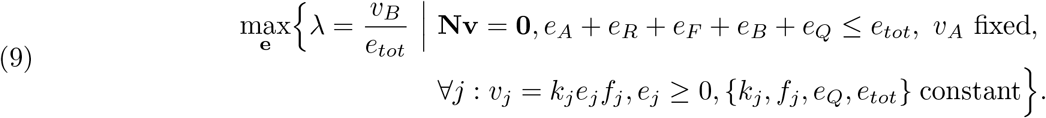

This includes an extra constraint on the carbon assimilation flux (*v*_*A*_).

Another way to realise carbon-limited growth is through chemostat cultivation. In this setup, the growth rate is set by the experimentalist through the dilution rate. The cells in the culture that survive are the ones that are able to minimise their carbon uptake flux to sustain this growth rate [45]. This also limits the concentration of the carbon source in the chemostat. The optimisation problem is thus formulated as one in which the assimilation rate of carbon is to be minimised per unit of biomass produced,

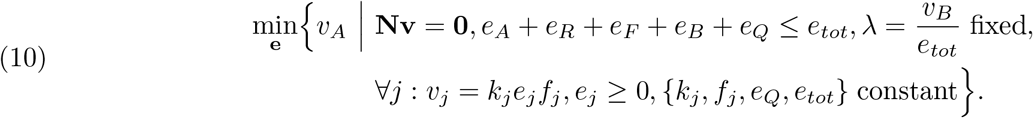

Mathematically, the optimisation problems (9) and (10) are equivalent. In both formulations, we are in the end maximising the ratio 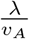, which is proportional to the biomass yield per mole carbon source,

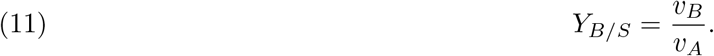

Both modes each have a different biomass yield. Since the fixed vectors of their corresponding EFMs (eq. (4)) are normalised with respect to the biosynthetic flux *v*_*B*_, their entries are yields. The biomass yield of each mode corresponds to the ratio of the last over the first entry of their EFM vector

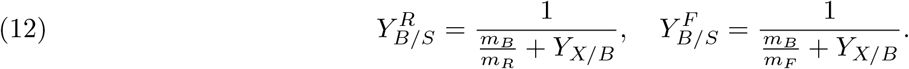

Because the *R* pathway is assumed to be most carbon-efficient (*m*_*R*_ *> m*_*F*_), the *R*-mode also has the highest yield: 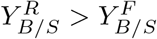 . It therefore requires the lowest assimilation flux to sustain a certain growth rate.

When multiple constraints become active, the *v*_*A*_-minimising flux vector can be a convex combination (8) of the EFMs. The optimal yield is now a convex combination of the yield of the two strategies,

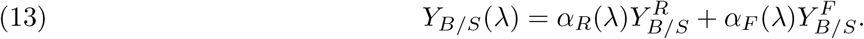

Clearly, the optimal yield is then always lower than or equal to the maximal yield 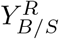.

We now show that if enzyme saturation levels are assumed to be constant (an assumption used in many proteome-constrained models [16; 20; 29; 34]), and cells have a constant total enzyme concentration across different conditions, we can identify a protein pool that is not related to growth but still needs to be made by the cell at low growth rates. As we will see, as the growth rate increases, this pool shrinks until it vanishes, and at this point, the cell needs to start using the carbon-inefficient pathway to grow even faster.

We start from the optimisation problem in eq. (10). Setting *λ* = *v*_*B*_*/e*_*tot*_ implicitly sets a constraint on *e*_*B*_, since *v*_*B*_ = *k*_*B*_*e*_*B*_*f*_*B*_, and *k*_*B*_ and *f*_*B*_ are assumed to be fixed,

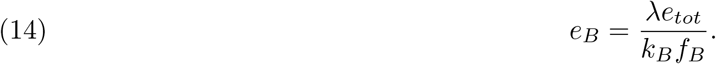

If the total protein constraint is not hit in the optimum, i.e., *e*_*A*_ + *e*_*R*_ + *e*_*F*_ + *e*_*B*_ + *e*_*Q*_ *< e*_*tot*_, then there exists a pool of enzymes not associated with growth,

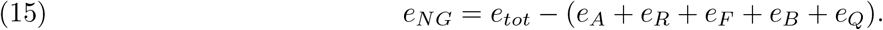

So depending on the growth rate, either 1 or 2 protein constraints are hit: certainly (14), and possibly

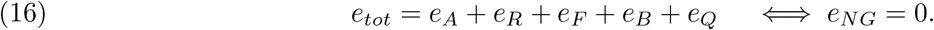

From previous work [38], we thus expect a switch from one EFM to a mix of two as the growth rate increases and the second constraint is hit. Other models that exhibit a shift between catabolic modes use a similar decomposition of the proteome in growth-associated and maintenance sectors [6; 20; 46] but typically without the *e*_*NG*_ pool we have identified here.

### 2.4 Reducing large genome-scale metabolic models to small core models

In Supplementary Section S8 we provide a protocol for deriving a core metabolic network from a large (genome-scale) network, based on the decomposition of the metabolic network in different functional processes as presented in Section 2.1. We derive equations for the (average) metabolic fluxes *v*_*j*_ for each process that are approximated in terms of averages of both the kinetic parameters and protein concentrations per sector. This allows for calibration of the core model with experimental data. To illustrate this, we apply this protocol to the yeast *S. cerevisiae* and derive a core model that describes its metabolic shift. Other reduction methods for large models have been derived by Erdrich et al. [47] and Hädicke and Klamt [48].

## 3 Results

### 3.1 The carbon-efficient regime of nutrient-limited growth: 0 *< λ < λ*_*c*_

Nutrient-limited growth can be attained in a chemostat [45] and by titration of the expression of the transporter of the nutrient [20; 44]. The nutrient we are considering is a source for both carbon and energy. We shall mostly be referring to aerobic growth on a sugar, as this is the best understood condition for the onset of overflow metabolism. The growth rate *λ* rises as function of the dilution rate in the chemostat and the titrated level of the transporter from a value close to 0 to the maximal growth rate *λ*_*max*_. When overflow occurs it does so at an intermediate critical growth rate *λ*_*c*_. What all these cases have in common is a shift at *λ* = *λ*_*c*_ from a carbon-efficient mode of metabolism (i.e., a high yield of biomass on the carbon source; respiration) to a less carbon-efficient mode of metabolism (overflow metabolism; fermentation). In all these cases, the carbon-efficient mode of metabolism, active exclusively below *λ*_*c*_, synthesizes more ATP per unit carbon source than the carbon-inefficient mode, which gradually overtakes the carbon-efficient mode in the regime *λ*_*c*_ *< λ < λ*_*max*_.

We investigate the onset of overflow in two experimental conditions. First, we focus on a carbon-limited chemostat. After that, we consider the shift as function of the growth rate on different carbon sources in carbon uptake-limited batch cultures.

One question we need to answer is why the carbon-efficient mode is favored over the inefficient mode in regime 0 *< λ < λ*_*c*_ and why the efficient mode cannot be used exclusively for higher growth rates. This becomes clear from the optimisation problem formulation of the chemostat (eq. (10)). The carbon assimilation rate *v*_*A*_ is minimised given a set growth rate (equal to the dilution rate of the chemostat). The set growth rate implies that we set the protein pool of biomass formation *e*_*B*_ equal to 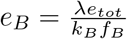. This is the only active protein expression constraint in the regime 0 *< λ < λ*_*c*_. Accordingly, a single EFM is active. Since in this case the optimisation problem is equivalent to maximising the yield of biomass on the carbon source *Y*_*B/S*_ = *v*_*B*_*/v*_*A*_, the carbon-efficient mode is the optimal solution because 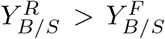 (eq. (12)). This explains why the carbon-inefficient mode of metabolism is not used in the growth rate regime 0 *< λ < λ*_*c*_.

In this growth rate regime, the carbon-efficient pathway may therefore be studied in isolation by setting the protein pool of the carbon-inefficient mode to zero (*e*_*F*_ = 0). The steady-state equations (1) and (5) now have a unique solution for the protein pools as function of the (set) growth rate. Since it is customary in the field to use protein fractions *ϕ*, i.e. protein concentrations normalised to the total protein concentration, we define, e.g., *ϕ*_*A*_ = *e*_*A*_*/e*_*tot*_.

In terms of protein fractions, the steady-state solution of the carbon-efficient, *R*-strategy is given by

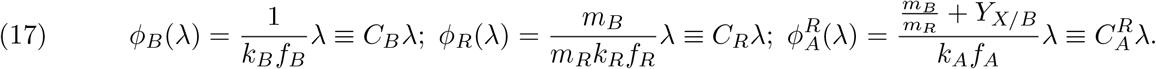

Here we introduced the constants^2^ *C*_*j*_ to collect corresponding parameter combinations. These constants may be interpreted as (lumped) protein costs for each lumped reaction. These equations indicate that the protein fraction are proportional to the growth rate, which is in agreement with experimental data [29; 49]).

Since *e*_*tot*_ is fixed, the increase in the growth-associated protein is accompanied by a proportional decrease of the growth-unassociated protein pool. Since *e*_*Q*_ is assumed fixed, this decreasing pool has to be *e*_*NG*_, and

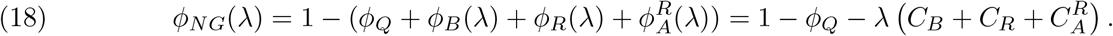

This behavior is also found in experimental data of protein sectors as function of the growth rate [11]. The last equation indicates the decrease with growth rate of the growth-unassociated protein pool and results from substituting the protein fraction relations in the protein fraction conservation equation.

The region in Figure 2A up to the critical growth rate, indicated by the dashed line, presents the protein fractions of the efficient strategy as function of the growth rate. The same region in Figure 2B presents the biomass yield, which is constant and maximal, and thus gives minimal carbon assimilation (rate).

**Figure 2.**
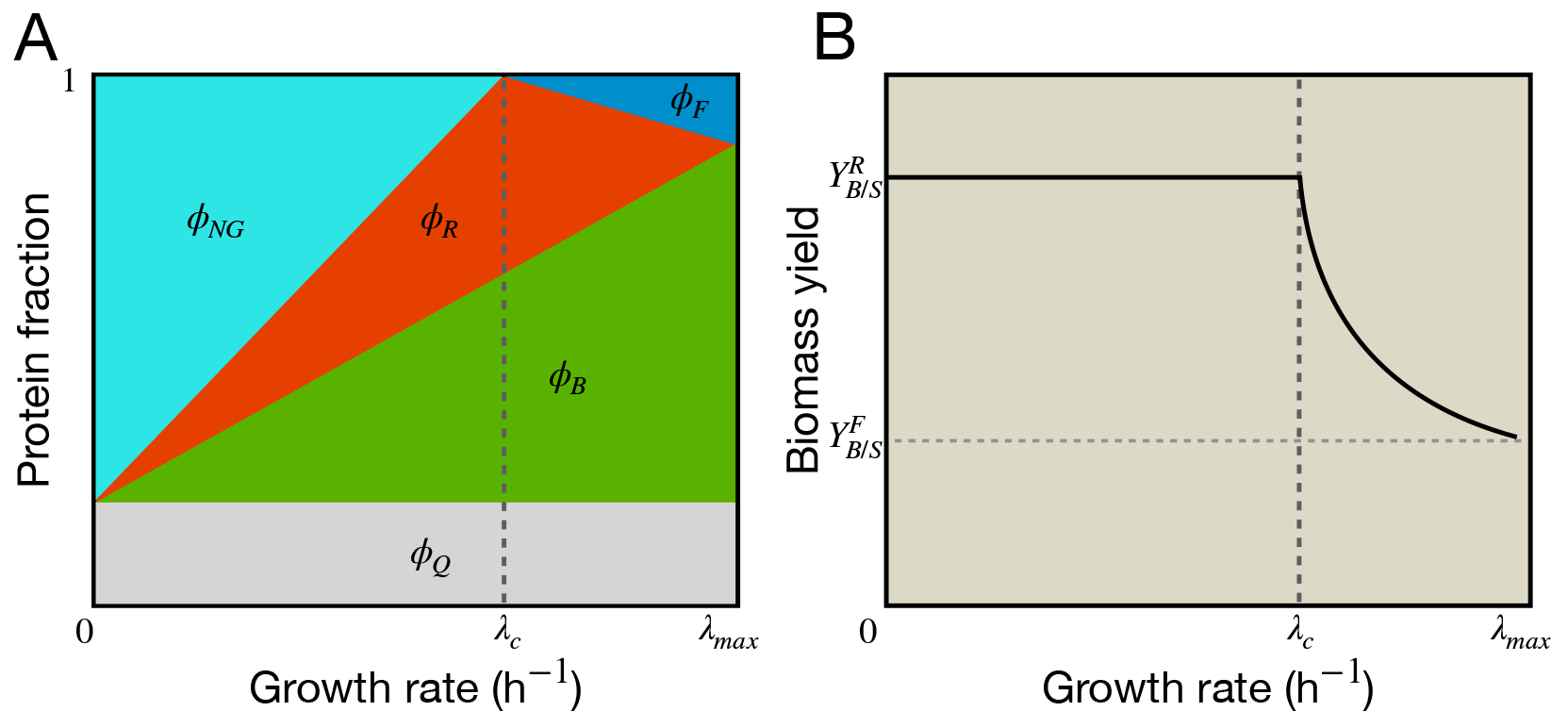
A shift from the carbon-efficient mode to the inefficient mode occurs when all available protein is invested in growth. A) Growth-associated protein fractions (*ϕ* _*R*_, *ϕ*_*B*_) increase linearly with the growth rate, at the expense of the *NG*-sector. Beyond the critical growth rate, the *R*-sector is replaced by the (smaller) *F* -sector. The assimilation sector (*ϕ* _*A*_) is omitted for clarity. B) The biomass yield is constant and maximal when only the carbon-efficient mode is active. Beyond the critical growth rate, the yield decreases because the carbon-inefficient mode has a lower yield.

The cell is able to increase its growth rate using only the carbon-efficient mode until all protein available for growth, i.e., *e*_*tot*_ *— e*_*Q*_ has been invested. The maximum is thus reached when the growth-unassociated protein sector *e*_*NG*_ is depleted. The cell has now reached its *critical growth rate λ*_*c*_. Solving eq. (18) at *ϕ*_*NG*_ = 0 for *λ*, we find

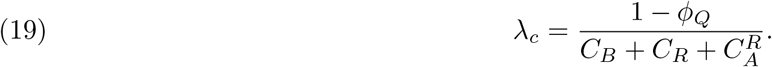

The depletion of the growth-unassociated protein pool constituted the second protein constraint that is hit, in addition to the one that was throughout the regime 0 *< λ < λ*_*c*_, i.e.,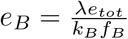 The growth rate can now only increase further if a part of the protein spent on the carbon-efficient mode is reinvested into another mode that is less carbon-efficient, but is more proteome-efficient, and into anabolism. In terms of the EFM theory [38], a second EFM mixes in with a first when a second protein expression constraint becomes active provided that the second strategy has the required behavior. In the next section, we will identify under which conditions this can occur.

Finally, we note that equation (18) when solved for *λ*, i.e.,

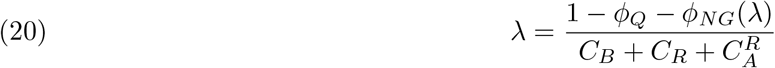

captures the trade-off between the instantaneous growth rate *λ* and the fraction of protein devoted to growth-unassociated processes [8]. These are often alternative nutrient uptake systems and stress tolerance [11]. The equation thus indicates that a higher investment in preparation for possible future conditions is concomitant with a decrease in the instantaneous growth rate. Put differently, there is a trade-off between the instantaneous and long-term growth rate [8]. These insights agree with experimental data; *E. coli* samples taken from an aerobic, glucose-limited chemostat below *λ*_*c*_ proved more adaptive and stress tolerant than samples taken at growth rate close to the maximal growth rate (above *λ*_*c*_) [9].

### 3.2 The overflow regime (*λ*_*c*_ *< λ < λ*_*max*_) during nutrient-limited growth

In order for growth to continue at a rate above *λ*_*c*_, another energy-generating strategy is required. The growth rate is directly proportional to the concentration of protein *e*_*B*_ invested. With a fixed total concentration, and no more growth-unassociated protein to spend, the protein spent on the carbon-efficient pathway needs to be reallocated to the other mode. This second, carbon-inefficient strategy should have a higher ATP synthesis rate than the carbon-efficient mode, but require a lower protein fraction. In terms of the model, this means that the protein costs for ATP synthesis are lower for the carbon-inefficient mode than for the carbon-efficient mode. This can be achieved by the inefficient mode having a higher activity per unit protein and/or because the associated metabolism relies on fewer enzymes (e.g. glycolysis + fermentation enzymes *<* glycolysis + TCA cycle + respiratory chain enzymes). When all the protein *e*_*R*_ that was previously invested in the carbon-efficient mode is completely used and replaced by *e*_*F*_ to catalyse the carbon-inefficient mode exclusively, the maximal growth rate *λ*_*max*_ is reached. The cell now only displays the carbon-inefficient strategy.

In the *λ*_*c*_ *< λ < λ*_*max*_ regime, the optimisation problem (10) also admits an analytical solution for the protein fractions. This is derived in Supplementary Section S3 and agrees with Basan et al. [20]. In this growth rate regime, the optimal strategy is a convex combination (8) of both strategies. The corresponding convex coefficients *α*_*R*_(*λ*), *α*_*F*_ (*λ*) result in linear protein fractions *ϕ* _*j*_(*λ*). An increase in the convex coefficient *α*_*F*_ (*λ*) inherently decreases its counterpart *α*_*R*_(*λ*) as they add up to one, thereby reflecting the exchange of the two strategies. The resulting change in the protein fractions as function of the growth rate is depicted in Figure 2A, after the dashed line.

The carbon-inefficient mode synthesizes less ATP per glucose while the needed ATP supply rate increases linearly with growth rate. The expectation is thus that the carbon uptake rate in the chemostat rises more steeply for *λ > λ*_*c*_ than below it. This is indeed the case in such studies [2; 25] and Figure 5C. The slope of the glucose uptake rate (*v*_*A*_) as function of the growth rate (*λ*) equals the inverse of the biomass yield on glucose. Since this slope is higher above *λ*_*c*_ we conclude that the yield of the carbon-inefficient mode is less than that of the efficient mode, i.e. 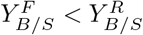 (as we assumed, cf. eq. (12)).

Since the optimisation still involves minimising the nutrient uptake for a given growth rate, we are still searching for the highest biomass yield on the carbon source, but now in accordance to two mixed strategies (eq.(13)). The resulting yield is therefore always below 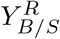 as the carbon-inefficient mode is mixed in. This is shown in Figure 2B after the dashed line. When the maximal growth rate is reached, the optimal yield equals 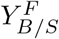.

From the above discussion, we can conclude that a necessary condition for the onset of overflow metabolism is for the second metabolic strategy to be carbon-inefficient (so that it is not favored as the main mode at low growth rates (*λ < λ*_*c*_)), but also that it is able to sustain a growth rate that exceeds the critical growth rate. In other words, *λ*_*max*_ *> λ*_*c*_.

Analogous to eq. (17)-(19), we study the carbon-inefficient mode in isolation to determine *λ*_*max*_. Setting *ϕ* _*R*_(*λ*) = 0 and solving the steady-state equations (1) and (5) for the remaining growth-associated protein fractions yields

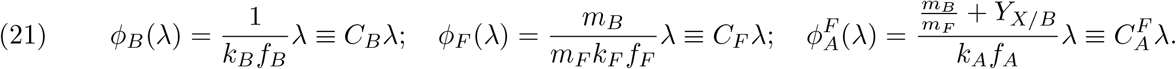

The maximal growth rate *λ*_*max*_ is found in exactly the same way as *λ*_*c*_ before, as explained in Supplementary Section S2. We find

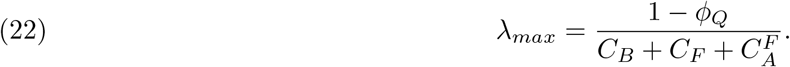

We immediately observe that *λ*_*c*_ *< λ*_*max*_ when 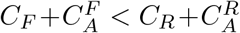 such that 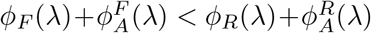 for all *λ*’s: the carbon-inefficient mode is more proteome-efficient.

Figure 2 may thus be summarised as follows. For growth rates above the critical growth rate, the (optimal) protein fraction *ϕ*_*R*_(*λ*) decreases until it is absent. At this point, the employed strategy has completely shifted^3^ from the efficient to the inefficient mode and the protein fractions are given by eq. (21). For the yield (13), this implies that 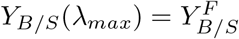. Since no alternative strategy exists that can further increase the growth rate, the maximal growth rate of the carbon-inefficient mode equals the absolute maximal growth rate.

### 3.3 The proteome efficiency of fermentation versus respiration as an overflow condition

In the previous section, we concluded that the carbon-inefficient catabolic mode can take over from the carbon-efficient catabolic mode above *λ*_*c*_ when the first has a maximal growth rate that exceeds the maximal growth rate of the second. This higher maximal growth rate is achieved at a lower protein fraction that is allocated to catabolism, since at higher growth rate biomass formation requires more protein. Hence, per unit protein the carbon-inefficient mode needs to achieve a higher ATP synthesis rate than the carbon-efficient mode. This insight underlies the introduction of the concept of proteome efficiency by others [16; 20; 29; 34]. It is loosely defined as the ATP synthesis rate per unit of protein and denoted by *ϵ*. In our model we define it by dividing by the total protein concentration expended to sustain the flux *v*_*R*_ and *v*_*F*_, respectively,

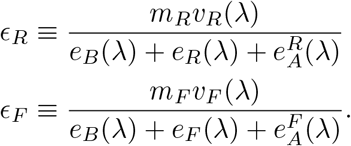

Using eq. (17), (21) and the steady-state equations (1) and (5), one may simplify these to

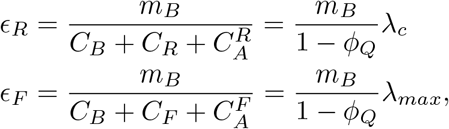

showing that these proteome efficiencies are directly related to the respective maximal growth rates. Explaining the shift in terms of the maximal growth rates or the proteome efficiencies of the strategies are therefore equivalent. In Supplementary Sections S4 and S16, we examine in detail the different definitions for the proteome efficiency. We show that, as long as protein costs are properly accounted for, they all lead to the same result: a shift to the carbon-inefficient mode caused by a constraint on the total protein content implies its higher proteome efficiency.

### 3.4 The critical and maximal growth rate respond similarly to physiological perturbations that affect both metabolic modes

Both the critical (eq. (19)) and maximal growth rate (eq. (22)) are proportional to 1 *— ϕ*_*Q*_. This shows that varying the Q-pool by overexpressing a protein that does not contribute to growth is expected to shift both the critical and maximal growth rate with the same factor. Treatment of cells with antibiotics gives a similar physiological perturbation of both modes by inhibiting translation. This corresponds to the decrease of the catalytic constant of biomass synthesis (*k*_*B*_) in the model, which increases the biosynthetic protein costs (*C*_*B*_). However, in contrast to the change in *ϕ*_*Q*_, this perturbation decreases the maximal growth rate by a larger factor than the critical growth rate. This indicates a larger effect on the carbon-inefficient mode due to its lower protein costs. Both these effects were indeed observed by Basan et al. [20]. Here, we formulate these in terms of biochemically interpretable parameters.

Finally, we note that the critical and maximal growth rate both depend on the *C*-coefficients representing the lumped protein costs of the growth-associated processes (each resembling a lumped metabolic pathway). All the *C*-coefficients (eq. (17) and (21)) are inversely proportional to the enzyme saturation degrees of the lumped pathways. These relations imply that both the critical and the maximal growth rate increase with the enzyme saturation degree. Less enzyme is then needed to reach a certain process flux and therefore higher rates can be reached.

### 3.5 Physiological parameters that influence the occurrence of overflow metabolism

In the previous section we considered physiological perturbations that change both metabolic modes. Although these affect the onset of overflow metabolism, this will likely not alter its manifestation. This can only be altered when properties of one of the modes change. For instance, the P/O ratio of the respiratory chain may vary between species or varied for one species by perturbation. This parameter is directly related to the *m*_*R*_ parameter in the core model.

To be able to address how physiological parameters influence the occurrence of overflow metabolism, we consider the following dimensionless ratio of the maximal growth rate of the inefficient over the efficient mode,

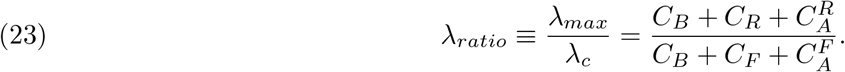

As concluded above, a shift occurs if *λ*_*max*_ *> λ*_*c*_; hence, when *λ*_*ratio*_ *>* 1. In general, the critical growth rate increases when the protein costs of the efficient mode decrease. This reduces *λ*_*ratio*_. Increasing the protein costs of the inefficient strategy causes a similar effect, since this reduces *λ*_*max*_. When *λ*_*ratio*_ ≤ 1 a metabolic shift will not occur.

We shall refer to *λ*_*ratio*_ = 1 as the break-even condition. Using eq. (23), this condition is written explicitly in terms of protein costs as

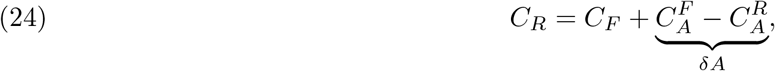

where *δA* is the difference in (carbon) assimilation costs for both strategies. This difference is generally positive, as the inefficient mode requires more carbon due to its lower yield.

The break-even condition separates the regime where a metabolic shift will occur (in blue) from the regime without a shift (in brown) in Figure 3A. This figure may therefore be interpreted as a phase diagram, similar to those used in physics. An organism is represented by its protein costs as a coordinate 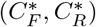.

**Figure 3.**
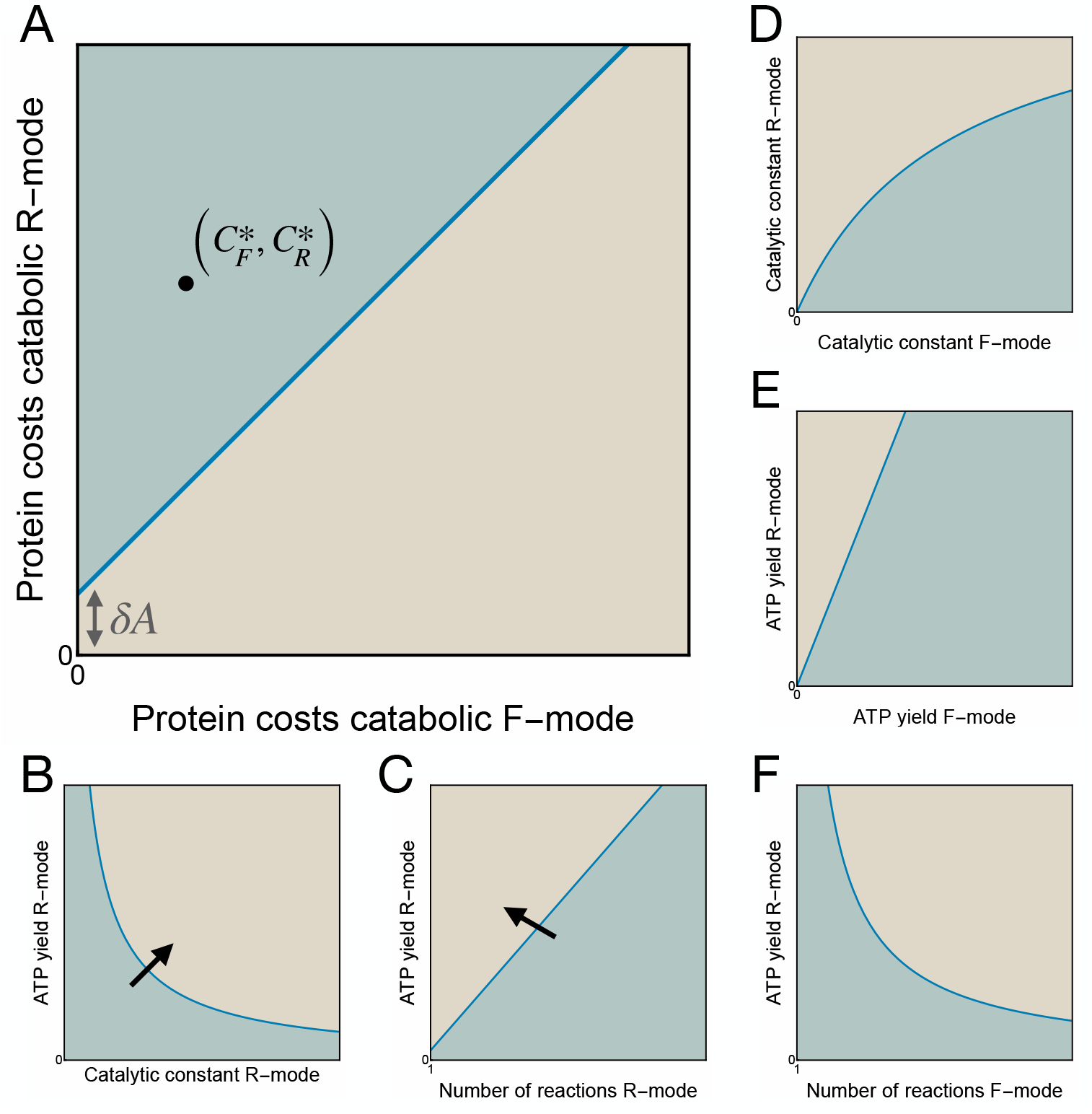
Parameter perturbations affect the occurrence of overflow metabolism. The break-even analysis in terms of A) catabolic protein costs and B)-F) several model parameters. For parameter configurations in the blue region, a metabolic shift occurs. In the brown region a shift will never occur. The arrow in B) and C) indicates the direction in which the break-even condition moves if the protein costs of the carbon-inefficient mode decrease.

To study this break-even condition for different parameters, it is useful to unpack a little how a large metabolic model may be condensed into a core model. Each reaction in the core model gives an average reaction rate. As detailed in Supplementary Section S8, expressions for these average rates include the number of individual enzymatic reactions in each lumped reaction. We denote these lengths by *N*_*A*_ for the number of reactions in assimilation, etc.

With these length parameters, the break-even condition can be written explicitly in terms of model parameters as

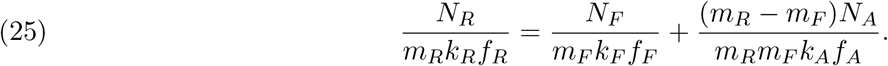

This follows from using the definitions of *C*_*j*_ from eq. (17) and (21).

The break-even condition allows us to investigate the dependence of two parameters (assuming all other parameters remain fixed). We generally find a curve that divides the two-parameter plane in regions. To illustrate this, Figures 3B-F show the (conceptual) phase diagrams for five different parameter pairs that are perturbed. All of these figures have a similar intuitive interpretation: only for values of the parameters that result in *C*_*R*_ *> C*_*F*_ + *δA*, a shift occurs.

For instance, Figure 3B shows that there is a trade-off between the ATP yield *m*_*R*_ and the catalytic constant *k*_*R*_. The effect of a change in the catalytic constants *k*_*R*_, *k*_*F*_ of both strategies on the metabolic shift is depicted in Figure 3D. Figure 3F shows that for increasing length *N*_*F*_ of the inefficient mode the protein costs *C*_*F*_ increase, so also the ATP yield *m*_*R*_ should decrease for a shift to occur. Additional examples for different parameters are given in Supplementary Section S5.

This break-even analysis may be used to clarify why some organisms or strains shift between metabolic strategies, but others do not [34]. It may be that *λ*_*ratio*_ *<* 1. This means that the carbon-efficient mode also has a higher proteome efficiency than the inefficient mode, so the inefficient mode can not reach growth rates exceeding *λ*_*c*_. A different reason for the absence of a shift is that the organism never attains its critical growth rate, and therefore the growth-unassociated protein sector never gets depleted.

### 3.6 The metabolic shift as function of the growth rate in (carbon uptake-limited) batch conditions with different carbon sources

In nutrient excess conditions, microbes attain their maximal growth rates. The associated growth rate maximisation problem is given in eq. (6). It follows from the previous sections that the growth rate is maximal when the growth-unassociated protein pool is empty (*e*_*NG*_ = 0). In nutrient excess conditions, neither the growth rate nor the assimilation rate is fixed. Therefore, there is only one active (proteome) constraint limiting growth in this case, and accordingly only a single EFM is active. This will be the EFM with the lowest protein costs. We therefore predict that the cell uses the carbon-inefficient mode in nutrient excess conditions.

As explained in Section 2.3, during carbon uptake-limited batch cultivation, the cell is limited by the assimilation rate *v*_*A*_ of carbon. Experimentally, this is achieved through titration of the carbon uptake system [20; 44]. This sets *e*_*A*_, and thereby the assimilation rate rate *v*_*A*_. The resulting optimisation problem (eq. (9)) is equivalent to the one in eq. (10) that describes chemostat conditions. Their solution in terms of optimal protein fractions is the same, but for eq. (9) it is parameterized in terms of *v*_*A*_ instead of *λ*. Therefore also in uptake-limited conditions, the efficient mode is optimal below a critical uptake rate

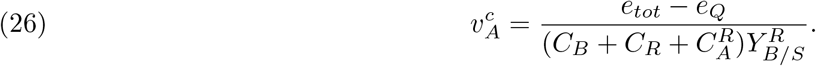

For higher carbon uptake rates, the growth-unassociated protein pool is empty and the carbon-inefficient strategy will occur. Due to the equivalence between the optimisations for chemostat and uptake-limited batch conditions, we find that 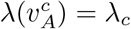.

Another way to realize uptake limitation experimentally is by varying the carbon source (at a constant titrated or wild type level of the uptake system). For each carbon source, the catalytic rate constant *k*_*A*_ of the assimilation pathway is different and represents the quality of the nutrient [5; 50].

Fixing *k*_*A*_ while also fixing the assimilation enzyme concentration *e*_*A*_ again results in a constraint on *v*_*A*_ that limits growth. However, there is a small but important difference in this implementation of uptake limitation with respect to the previous setting. When, instead, the carbon source is fixed (by the experimentalist), uptake limitation is only realized for a constant *e*_*A*_. Otherwise, the cell may increase this enzyme concentration at low-quality nutrients to alleviate the uptake limitation that constrains the growth rate. Whether it will do so depends on its gene-regulatory strategy. This minor distinction leads to different quantitative results.

The solution to this optimisation problem is now parameterized in terms of the nutrient quality *k*_*A*_. The shift from the efficient to the inefficient mode now initiates at a critical nutrient quality

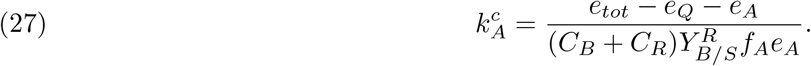

It depends on the constant value of *e*_*A*_ in two ways. This critical nutrient quality corresponds to a critical assimilation rate

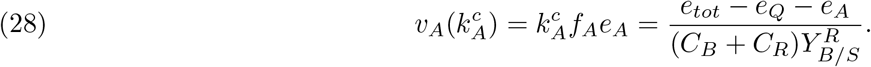

Due to the constant *e*_*A*_, the assimilation contribution appears as *e*_*A*_ in the numerator, rather than as the costs 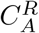 in the denominator in eq. (26) for 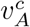 . The corresponding critical growth rate

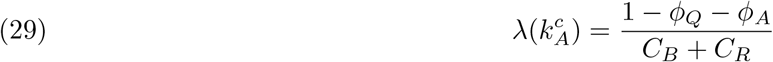

differs in the same way from the expression

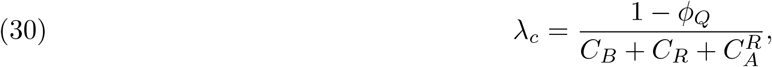

derived in Section 3.1. The analysis of the optimisation problem with *k*_*A*_ and *e*_*A*_ fixed is presented in more detail in Supplementary Section S6.

Our analysis of the difference between chemostat settings and varying nutrient quality in batch conditions allows us to understand why the patterns both display are so alike. It is consistent with the experimental data for *E. coli* obtained by Basan et al. [20]. They studied carbon uptake-limited cultivation of *E. coli* by varying titrated levels of the carbon uptake system on different glycolytic and non-glycolytic carbon sources. Both for increasing uptake titration and increasing nutrient quality, the authors observed an increase in the acetate production flux with the growth rate after a critical growth rate. This indicates a higher activity of the acetate-formation pathway, which is the more proteome-efficient pathway in *E. coli* [20].

What is particularly noteworthy is that the data seems to follow the same linear increase in the acetate production flux with the growth rate, both for increasing uptake titration and increasing nutrient quality. This ‘acetate line’, as it is called by the authors, is in the core model represented by the (overflow) flux *v*_*F*_ (*λ*). In Supplementary Section S6 we show that this flux, when parameterized by the nutrient quality *k*_*A*_, is a linear function of the growth rate, with both the slope and intersection with the *λ*-axis determined by the critical growth rate (29). Nevertheless, the critical growth rate 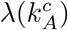 depends on both the constant assimilation fraction *ϕ*_*A*_ and the protein costs *C*_*j*_, which can differ per carbon source. Therefore, the linearity of *v*_*F*_ (*λ*(*k*_*A*_)) alone does not explain the observation of a single overflow flux line.

Proteomics data [46; 51] shows, however, that the assimilation fraction is generally small relative to other protein fractions. Furthermore, for glycolytic carbon sources, the cell uses similar catabolic and anabolic pathways, resulting in comparable protein costs *C*_*j*_. Together, this results in a value for the critical growth rate that is similar both for (small) values of *ϕ*_*A*_ and for different glycolytic carbon sources. In fact, small *ϕ*_*A*_ is equivalent to low assimilation costs 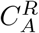 (with respect to other protein costs). In this limit, the different equations (29) and (30) for the critical growth rate reduce to the same expression. Consequently, this results in (approximately) one overflow flux line *v*_*F*_ (*λ*(*k*_*A*_)), because this is completely determined by the critical growth rate. This explains why the *E. coli* data from Basan et al. [20] for different experimental (carbon) uptake-limiting conditions seem to follow the same acetate line.

We can now also understand why the data for non-glycolytic carbon sources (see Extended Data Figure 1f in [20]) do *not* follow the acetate line. This is because the employed catabolic (and anabolic) pathways change for these carbon sources, thereby resulting in different associated protein costs and thus different values for the critical growth rate (29).

### 3.7 Characteristics of overflow metabolism in yeast are captured by a core model variant that follows from a genome-scale model

As explained in Section 2, large complex models used to analyse metabolic shifts, such as GEMs, may be reduced to a small core model, provided that equivalent constraints and degrees of freedom are taken into account. In this section, we illustrate this by adapting the core model to describe overflow metabolism in *S. cerevisiae* occurring at high growth rates in aerobic glucose-limited chemostat cultures. The reaction network corresponding to the yeast model is depicted in Figure 4. It is similar to the main core model sketched in Figure 1, but instead of combining carbon transport and phosphorylation into an assimilation pathway *A*, the yeast model describes this as separated carbon transport *T* with flux *v*_*T*_, and glycolysis *G* with flux *v*_*G*_ and pyruvate as its product. Furthermore, we assume the biosynthetic pathway *B* does not require any carbon in the yeast model. The analysis of this model focuses on the analogue of optimisation problem eq. (10).

**Figure 4.**
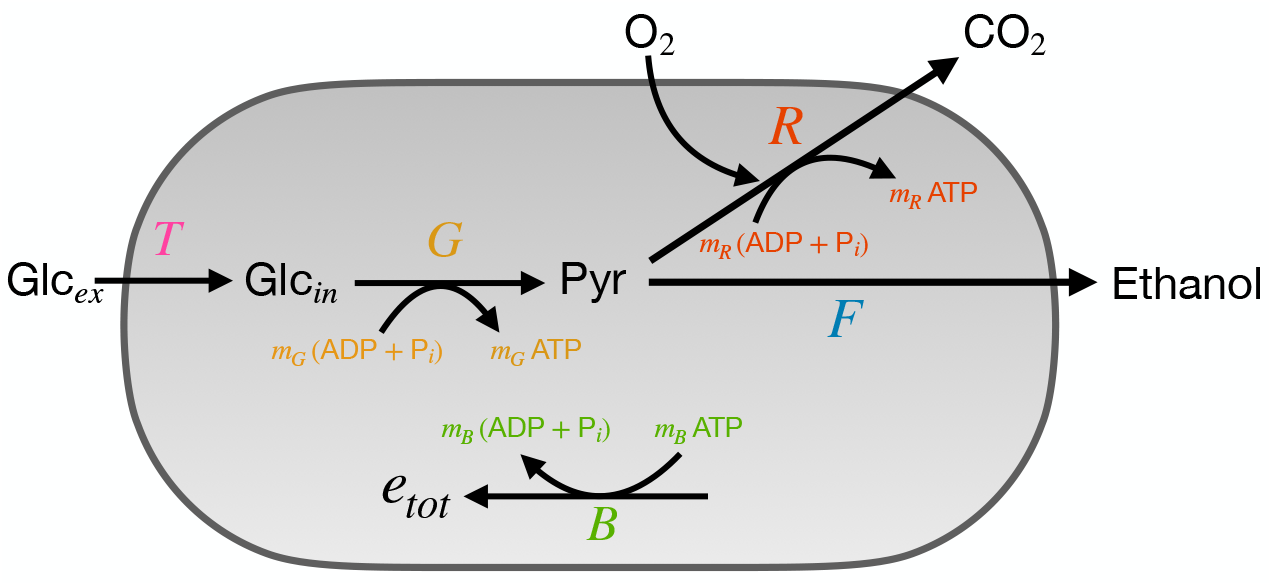
The metabolic network described by the yeast model. Glucose is imported by a transport reaction (*T*). Glycolysis (*G*) converts glucose into pyruvate, yielding 2 ATP per glucose. Pyruvate is further degraded into carbon dioxide by respiration (*R*) or into ethanol by fermentation (*F*). Respiration yields an additional 16 ATP per glucose. The generated ATP is used for biomass synthesis (*B*).

In Supplementary Section S8 we show that the rate *v*_*j*_ of lumped reaction *j* in the core model may be approximated in terms of averaged kinetic properties of all reactions in “sector” *j* as

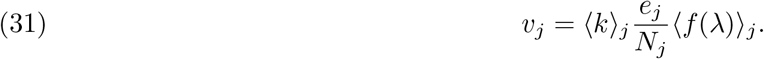

The rate *v*_*j*_ should then be interpreted as the mean rate of the reactions in sector *j*. Here, ⟨*k*⟩_*j*_ is the mean catalytic rate constant of these reactions. The enzyme concentration *e*_*j*_ in (31) signifies the total concentration of all enzymes involved in sector *j*, distributed among the *N*_*j*_ (metabolically distinct) reactions. Mean saturation levels ⟨*f* (*λ*)⟩_*j*_ that change with the growth rate are inferred from proteomics data from Elsemman et al. [29]. Other specific modifications to the core model for yeast, such as which pathways are included in each sector, are detailed in Supplementary Sections S7-S10.

We calibrate the model parameters to ensure that the critical and maximal growth rates are consistent with those observed during experimental cultivation of yeast. In the chemostat, ethanol formation starts at the critical dilution rate 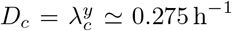 . The maximal growth rate as defined in eq. (22) should correspond to the point where yeast grows fully fermentative. However, this state is never reached in the chemostat. Therefore, we set the maximal growth rate of yeast to its growth rate during batch cultivation with excess glucose, which is 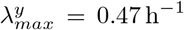 for the strain examined by Elsemman et al. [29]. A Mathematica implementation of the core model adapted to yeast is provided in the Supporting Information.

The optimal protein fractions obtained with the yeast model are depicted in Figure 5A as function of the growth rate. They exhibit dynamics qualitatively similar to Figure 2A, and may be interpreted analogously. Once the non-growth protein fraction is depleted at *D* = *D*_*c*_, an extra constraint becomes active and the model shifts from high-yield respiration to low-yield ethanol fermentation. The nonlinear relationship between the protein fractions and growth rate results from the saturations that now also change with the growth rate. Figure 5B shows qualitative agreement with the categorised proteomics data from Elsemman et al. [29]. The proteomics data show the presence of fermentative protein before the critical growth rate. This may be explained by preparatory expression of these enzymes. For example, the enzyme pyruvate decarboxylase is allosterically regulated and can therefore remain in an inactive state [52; 53]. These results are discussed in more detail in Supplementary Section S13. In Supplementary Section S14, we show that including changing saturation gives a better quantitative match between the model and the experimental data than for constant saturations.

**Figure 5.**
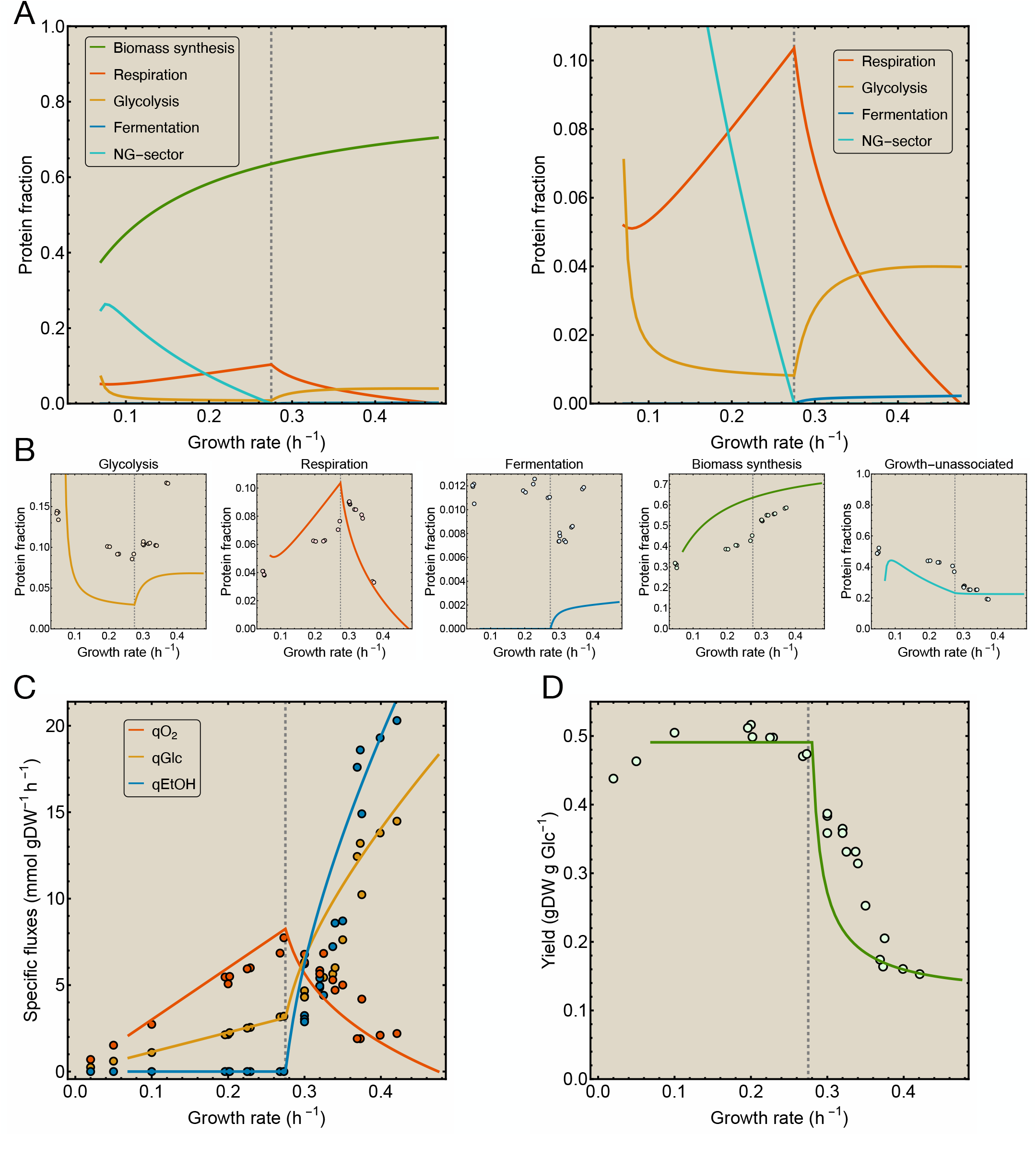
The yeast model captures the characteristics of overflow metabolism in yeast. A) Dynamics of the protein fractions as function of the growth rate. The second figure is enlarged to show the smaller fractions. B) The protein fractions predicted by the yeast model together with proteomics data [29]. C) Specific fluxes and D) Biomass yield predicted by the yeast model together with chemostat data [29]. In all figures, the dashed line indicates the critical growth rate. Data is reused with permission from [29] (Creative Commons Attribution 4.0 International License).

Specific uptake and production fluxes follow directly from normalising the model fluxes *v*_*j*_ by the dry weight density 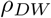, and accounting for the appropriate stoichiometric coefficient. The glucose uptake flux 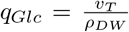 predicted by the model is given by the orange line in Figure 5C. Its faster increase after the critical growth rate results from the higher carbon requirement of fermentation. The oxygen uptake flux is 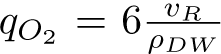, as respiration requires six moles of oxygen for the complete oxidation of one mole of glucose to carbon dioxide and water. It declines after the critical growth rate due to the replacement of respiration by fermentation, as the red line in Figure 5C shows. During fermentation, yeast metabolizes glucose into two ethanol, carbon dioxide and water. Thus, the ethanol production flux follows from the fermentation flux in the model as 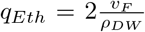. Ethanol excretion, shown by the blue line in Figure 5C, initiates after the critical growth rate. We compute the biomass yield via a unit conversion of the model prediction (eq. (13)), resulting in the green line in Figure 5D. At low growth rates it equals the respiration yield. When the critical growth rate is exceeded, the yield declines to the fermentation yield.

We have had to make a number of adjustments to make (some of) the specific fluxes and yield fit the experimental data, because of assumptions used in the yeast model. The model predictions shown in Figure 5C and D are the results after these adjustments. These are detailed in the Supplementary Sections S11 and S12.

In Figure 6 we give an illustration of the result from Section 3.6 for parameters of the yeast model and constant saturation factors. It shows that, during uptake-limited growth on different carbon sources, there is approximately one ethanol flux line when the transporter fraction (*ϕ* _*T*_) is small. Whether this also applies to yeast remains uncertain and requires experimental results similar to *E. coli* found by Basan et al. [20].

**Figure 6.**
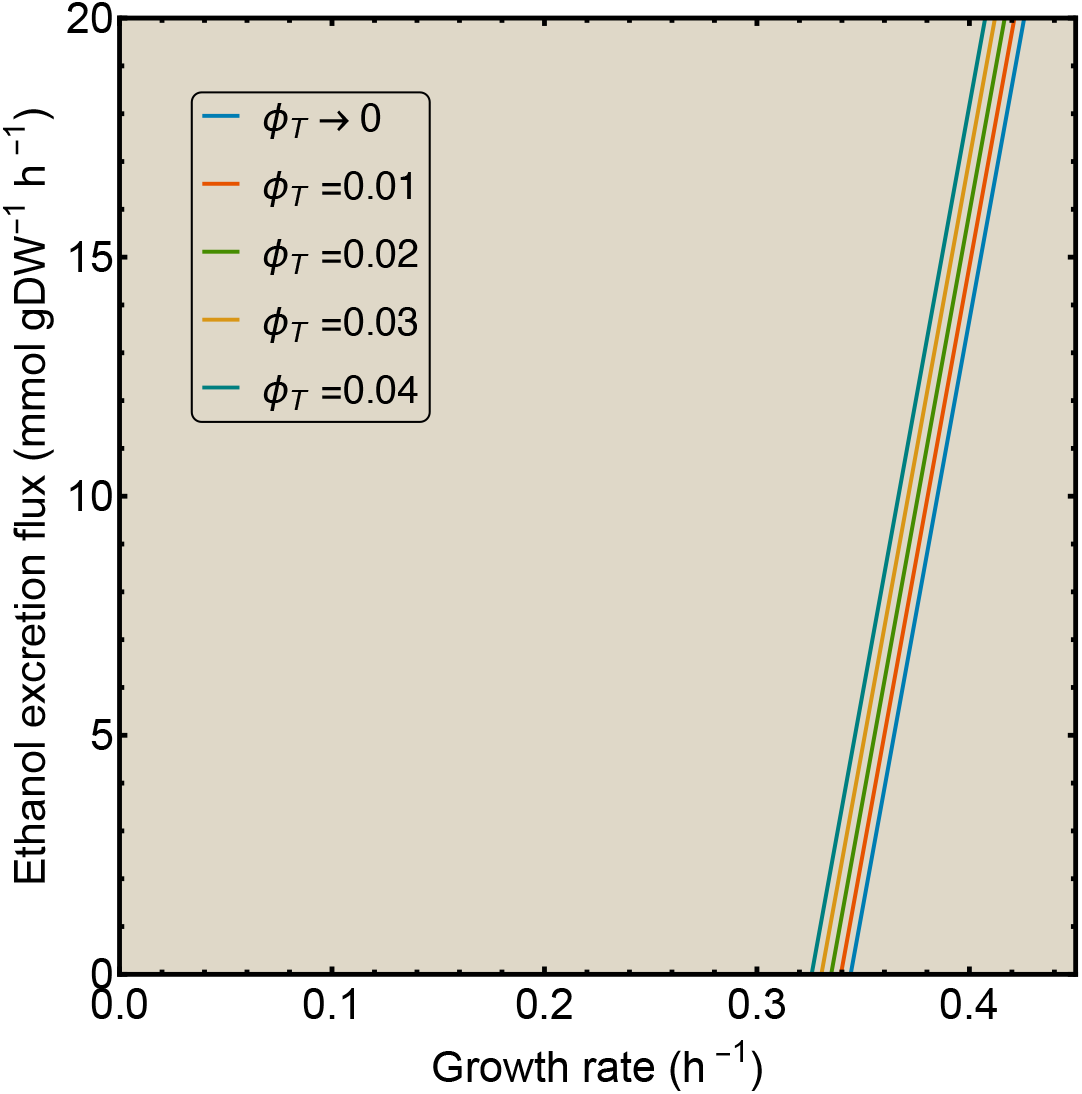
The metabolic shift is similar under chemostat and carbon uptake-limited batch conditions. Ethanol excretion flux as function of the growth rate for different values of constant transporter fraction (*ϕ*_*T*_). When *ϕ* _*T*_ is small, the corresponding ethanol excretion lines are parallel and close to each other, seemingly resulting in only one line. A Mathematica implementation is provided in the Supporting Information.

In Supplementary Sections S15-S17 we again analyse the respiration and fermentation strategies independently, compute their proteome efficiencies, and perform a break-even analysis.

## 4 Discussion & Conclusion

We have studied the conditions for the occurrence of overflow metabolism, an inefficient mode of metabolism, seemingly wasting the carbon and energy source. We identified two growth rate regimes separated by a critical growth rate, beyond which overflow metabolism starts. The carbon-efficient mode is favored below this growth rate because it has the maximal biomass yield on the substrate. At the critical growth rate, all available biosynthetic resources are used; the growth rate can only increase further if the maximal yield strategy is traded in for a suboptimal yield (i.e., carbon-inefficient) mode that is sufficiently proteome-efficient. We have shown that this is equivalent to the maximal growth rate of the carbon-inefficient mode exceeding the critical growth rate, which is the maximal growth rate of the carbon-efficient mode.

These results are illustrated and formally derived using a core model of cellular metabolism, but equally apply to genome-scale models. Indeed, we have outlined how such large-scale models may be reduced to a coarse-grained model, and have given an illustration from *S. cerevisiae* that the agreement with experimental data is qualitatively the same. We have also shown that the overflow line appearing in both chemostat cultures and in carbon uptake-limited batch cultures in different nutrient conditions essentially have the same origin, and can be explained using the same principles. Thereby, we compare and unify these (apparently) different conditions for the first time.

Our model is an extension of the model proposed in [20]. The novelty of our model is that, by introducing a growth-unassociated protein pool, we can study the growth rate regime below the critical growth rate (0 *< λ < λ*_*c*_). Another key element is that we explicitly assume constant enzyme saturation of growth-associated enzymes at all growth rates. This induces linear relations between protein pools and growth rate (eq. (17) and (21)), which is in agreement with proteomics data [29; 42; 49]). Constant enzyme saturations imply that a low growth rate, due to nutrient-limitation or a low-quality nutrient, can be attained when not all protein is allocated to growth processes. This allows for the expression of growth-unassociated proteins, that may be used for adaptation to alternative nutrients and stresses [11]. This enhances the long-term fitness of the microbe. At the critical growth rate, this pool has been completely replaced by growth proteins, giving priority to the instantaneous growth rate over adaptation [8]. A higher enzyme saturation of growth-associated enzymes increases the size of the growth-unassociated protein pool, the critical growth rate and the maximal growth rate. It can therefore be postulated as an objective of evolutionary maximisation.

In reality, saturations of individual enzymatic reactions change (marginally) with growth rate. The results of the coarse-grained model for *S. cerevisiae* show that including these still leads to expression of growth-unassociated proteins at low growth rates. Others [28; 29] also relied on a similar pool to describe overflow metabolism in yeasts. Moreover, accounting for changing saturations results in a better fit to proteomics data (Supplementary Section S14).

For the carbon-inefficient mode to have a higher maximal growth rate than the carbon-efficient one, its proteome efficiency needs to be higher. This occurs when the carbon-inefficient mode uses a catabolic pathway with fewer reactions than the efficient mode and/or deploys enzymes that are per-enzyme more active. Indeed, fermentation pathways generally consist of fewer reactions than respiratory metabolism. In Supplementary Sections S4 and S16 we show that the carbon-inefficient mode has a higher proteome efficiency irrespective of the definition of proteome efficiency that is used (as long as protein costs are accounted for properly). Different definitions therefore lead to the same results regarding the metabolic shift. By associating the proteome efficiency directly with the maximal growth rates of the two strategies we solve the problem that a clear definition of proteome efficiency was lacking [34; 54]: rather than focusing on proteome efficiency and how it is measured or defined, one may consider the maximal growth rate the pathway is able to sustain.

The conditions required to observe a metabolic shift that we derived are not always met. Indeed, overflow metabolism does not occur in all microbes, and the exceptions are enlightening. Some yeasts, such as *Pichia kluyveri*, do not overflow at excess glucose under aerobic conditions, while others like *S. cerevisiae* do. Battjes et al. [34] show that this difference between these two yeasts results from a difference in respiratory stoichiometry. From our perspective, the ratio *λ*_*c*_*/λ*_*max*_ is in that case greater than 1: the putative overflow pathway simply can not attain higher growth rates than the carbon-efficient one (respiration). This explanation is thus not complementary, but, in fact, equivalent.

We applied our theory to different environmental conditions, such as chemostats and (uptake-limited) batch cultures, as summarised in Figure 7. This agrees with chemostat data for *S. cerevisiae* [29]. The theory also provides a new explanation why varying both titrated levels of the uptake system and nutrient quality of glycolytic carbon sources in batch conditions leads to a single overflow flux line for *E. coli* [20]. Our theory also explains why growth on non-glycolytic carbon sources does not lead to the same trend, illustrating that the result from Basan et al. [20] is not universal. This agrees with a study by Wang [55] that applies a more extensive modeling approach to analyse overflow metabolism as function of the nutrient quality on both types of carbon sources.

**Figure 7.**
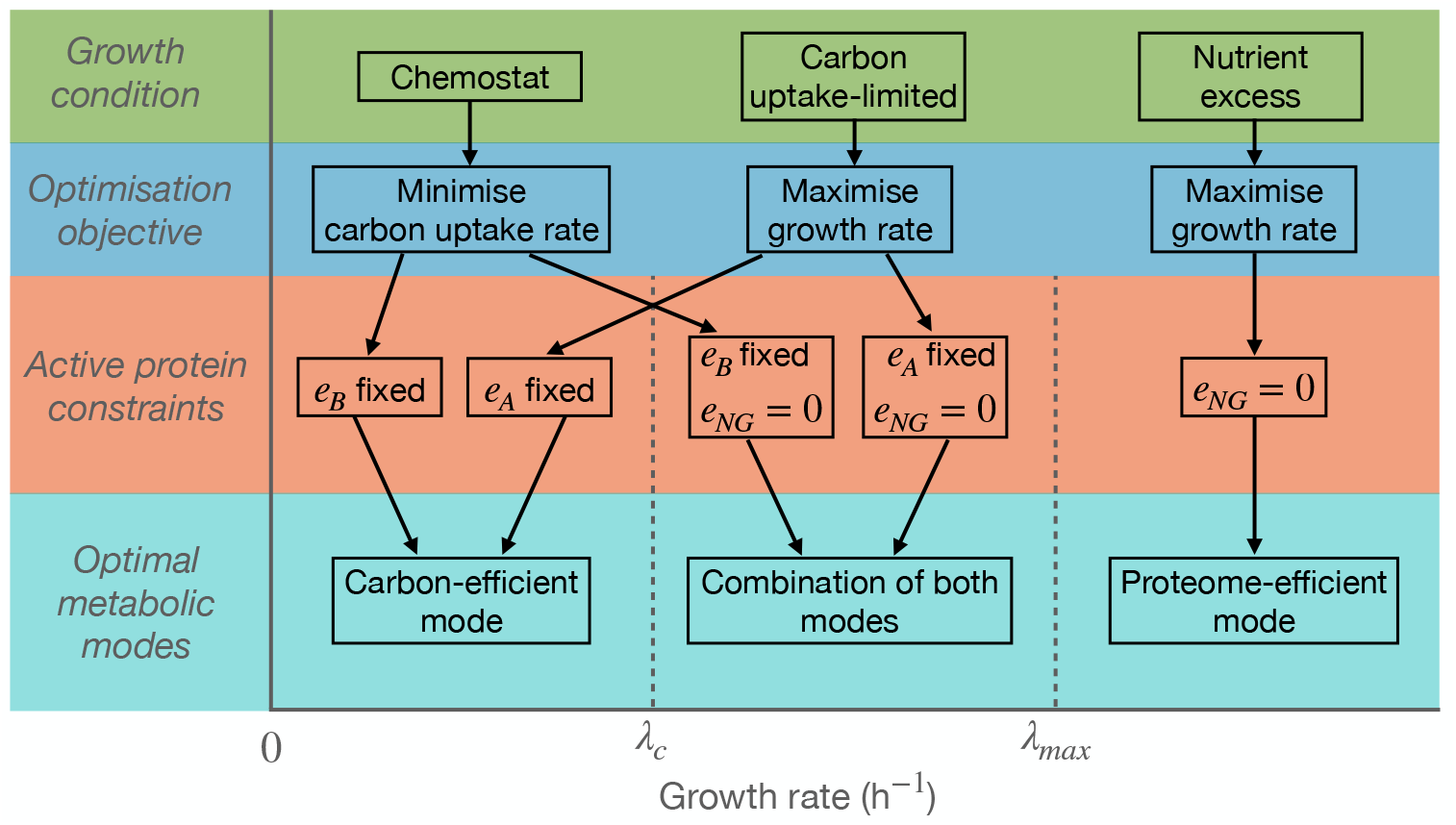
We predict optimal metabolic modes used in different experimental conditions with the core model.

Some interesting exceptions to our framework include growth of *S. cerevisiae* on trehalose and galactose in excess [29]. In carbon-excess conditions, we predict the use of the most proteome-efficient mode (fermentation), while the data shows that this yeast primarily respires on these carbon sources. For trehalose, we can understand this from its extracellular hydrolysis into glucose. This effectively causes carbon limitation, which clarifies the respiratory activity using our theory. A galactose-glucose trade-off [56; 57] may explain the use of respiration during growth on excess galactose.

The results of our core model may be expressed in terms of either the carbon uptake rate or the growth rate. We opted for the growth rate, as this is the controlled parameter during chemostat growth. However, the current version of the model is not suitable for analyzing the metabolic shift when different chemical-element sources limit growth, such as in Li et al. [58]. A different approach is also required to study metabolic shifts that do not seem to arise from proteome costs and constraints, such as in *L. lactis* [32]. Furthermore, the mean saturation functions, which we used in the coarse-grained model for *S. cerevisiae*, can still be improved and extended to better represent kinetic and thermodynamic factors [23; 59]. This may be useful when studying metabolic shifts that occur close to thermodynamic equilibrium [33].

Which mode of metabolism is favored, and how many are used, depends on the metabolic objective and the constraints that are limiting growth [38]. This reasoning is a key component in many models that describe shifts between metabolic strategies [6; 16; 20; 35; 36; 37; 38; 46; 55; 60; 61; 62]. Especially some of the coarse-grained models [6; 20; 38], which served as inspiration for the core model presented here, overlap in many aspects. This is inevitable, as they all aim to describe the same phenomenon with only a few ingredients.

We are thus by no means the first to study overflow metabolism. The goal of this work is *not* to elucidate which growth-limiting constraints are hit and to provide yet another explanation for this phenomenon [17]. Instead, the coarse-grained approach presented here is used to formulate the problem in general terms, so that different experimental conditions may be captured and thus unified. We have tried to bridge the gap between the large genome-scale and coarse-grained models, thereby deepening our understanding of shifts in metabolic strategies across different organisms and environmental conditions.

## Supporting Information

### Supplementary Appendices S1-S17

Supplementary sections with glossary, derivations, and additional background information on the core model and its adaptation to *S. cerevisiae*. (PDF)

### Source codes

Mathematica notebooks used for numerical implementations of the core model adapted to *S. cerevisiae*. These notebooks are used to create Figures 5, 6, S2, S3, S4, S5, S6 and S7, and compute the proteome efficiencies in Table S5. Source codes are available on GitHub (https://github.com/MaartenJDroste/Core-model-overflow-metabolism). (NB)

### Datasets

Excel files with (adjusted) data from chemostat experiments for *S. cerevisiae*, reused with permission from Elsemman et al. [29] (Creative Commons Attribution 4.0 International License). This data is used to calibrate the yeast model or for comparison of the model results to experimental data. Datasets are available on GitHub (https://github.com/MaartenJDroste/Core-model-overflow-metabolism). (XLSX)

## Acknowledgements

We thank Pranas Grigaitis for his input on the GEM-reduction and parameterization of the core model variant for yeast. We would like to thank our colleagues Julius Battjes, Maaike Remeijer, Jeroen van Kasteren and Bas Teusink for critical and stimulating discussions. MJD and RP acknowledge funding by NWO grant 613.009.131 ‘Control of maximal growth rate by single-celled organisms’.

## Footnotes

1 For glycolytic carbon sources, like glucose, the assimilation pathway consists of uptake and upper glycolysis. So, in these cases, the intermediate *X* represents DHAP and GA3P.

2 The saturation of the assimilation pathway (*f*_*A*_) is in fact not constant, but depends on the growth rate via the Monod relation. Then we find a unique solution for the protein pools only when an extra degree of freedom is added. Since the assimilation fraction is typically small, we expect the effect of a variable *f*_*A*_ to be negligible, so we set it constant.

3 In terms of the convex coefficients, *α*_*F*_ (*λ*_*max*_)=1 and *α*_*R*_(*λ*_*max*_)= 0.

