## Supplementary Appendices for "On the conditions for shifts in metabolic strategies"

### S1 Glossary

$C_j$: Protein costs associated with process $j$.
$D_c$: Critical dilution rate after which yeast shifts from respiration to ethanol fermentation in glucose-limited chemostats.
$e_j$: Concentration of the enzyme catalysing the lumped reaction that represents process $j$.
$e_{NG}$: The concentration of growth-unassociated protein that changes with growth rate.
$e_Q$: The concentration of growth-unassociated protein that is used for maintenance and other 'household' processes, assumed fixed across conditions.
$e_{tot}$: The total concentration of protein in a cell, assumed fixed across conditions.
$f_j$: Saturation factor of the enzyme catalysing the lumped reaction that represents process $j$.
$\langle f(\lambda)\rangle_j$: Saturation function that describes the mean enzyme saturation factor of the individual enzymatic reactions included in lumped reaction $j$ as a function of the growth rate in yeast.
$k_j$: Catalytic rate constant of the lumped reaction that represents process $j$.
$\langle k\rangle_j$: The mean catalytic rate constant of the individual enzymatic reactions included in lumped reaction $j$.
$k_A^c$: Critical catalytic rate constant of the assimilation pathway during carbon uptake-limited growth.
$m_B$: The amount of ATP required to synthesize one mole biomass.
$m_j$: ATP yield per mole of the carbon source, for reactions $j = R, F, G$.
$N_j$: Number of individual enzymatic reactions included in the sector represented by lumped reaction $j$.
$q$: Specific uptake or excretion rate of a compound per gram of dryweight biomass per hour.
S: Carbon source.
$v_j$: The rate of the lumped reaction that represents process $j$.
$v_A^c$: Critical assimilation rate during carbon uptake-limited growth.
$\mathbf{v}_F$: EFM corresponding to the carbon-inefficient (F) mode.
$\mathbf{v}_R$: EFM corresponding to the carbon-efficient (R) mode.
X: Carbon intermediate.
$Y_{B/S}$: Biomass yield per mole carbon source.
$Y_{X/B}$: The amount of carbon intermediate consumed to synthesize one mole of biomass.

$\alpha$: Convex coefficient associated with an EFM.
$\epsilon$: Proteome efficiency, i.e., the ATP synthesis rate per unit of protein.
$\lambda$: Growth rate of a microbial culture.
$\lambda_c$: Critical growth rate at which a shift to a different metabolic strategy commences.
$\lambda_c^y$: Critical growth rate of yeast during aerobic glucose-limited chemostat cultivation (equal to $D_c$).
$\lambda_{max}$: Maximal growth rate of a microbial culture.
$\lambda_{max}^y$: Maximal growth rate of yeast in batch conditions.
$\lambda_{ratio}$: The ratio of the maximal growth rate over the critical growth rate.
$\phi_j$: Fraction of the total proteome allocated to lumped reaction $j$.

### S2 Derivation of the equation for $\lambda_{max}$

To determine an equation for $\lambda_{max}$, we study the carbon-inefficient mode in isolation. Since $\phi_R = 0$, we solve the steady-state equations (1) and (5) for the remaining protein fractions, yielding

$$\text{(S1)} \qquad \phi_B(\lambda) = \frac{1}{k_B f_B}\lambda \equiv C_B\lambda; \quad \phi_F(\lambda) = \frac{m_B}{m_F k_F f_F}\lambda \equiv C_F\lambda; \quad \phi_A^F(\lambda) = \frac{\frac{m_B}{m_F} + Y_{X/B}}{k_A f_A}\lambda \equiv C_A^F\lambda.$$

The solution for the biosynthetic protein fraction $\phi_B$ matches its counterpart for the carbon-efficient mode (eq. (17)), demonstrating that biomass synthesis is completely independent of the used catabolic mode in this model.

The growth-associated protein fractions in eq. (S1) are also proportional to the growth rate. The remaining protein fraction is used for growth-unassociated tasks. In other words, when the carbon-inefficient mode is employed in isolation, it has its own growth-unassociated fraction $\phi_Q + \phi_{NG}^F$. This fraction decreases with growth rate because the total protein concentration is fixed. Since $e_Q$ is assumed fixed, this decreasing pool is again $e_{NG}^F$, with corresponding fraction

$$\text{(S2)} \qquad \phi_{NG}^F(\lambda) = 1 - (\phi_Q + \phi_B(\lambda) + \phi_F(\lambda) + \phi_A^F(\lambda)) = 1 - \phi_Q - \lambda\left(C_B + C_F + C_A^F\right).$$

Similar to the carbon-efficient mode, the carbon-inefficient mode reaches its maximal growth rate when the growth-unassociated protein sector $e_{NG}^F$ is depleted. Solving eq. (S2) at $\phi_{NG}^F = 0$ for $\lambda$ gives

$$\text{(S3)} \qquad \lambda_{max} = \frac{1 - \phi_Q}{C_B + C_F + C_A^F}.$$

When this growth rate is reached, no alternative strategy exists to further increase the growth rate. So, the maximal growth rate of the carbon-inefficient mode is in fact the maximal growth rate of the cell in this model.

### S3 Analytical solution of the core model in the regime $\lambda_c < \lambda < \lambda_{max}$

The optimisation problem

$$\text{(S4)} \qquad \begin{aligned} \min_{\mathbf{e}} \Big\{ v_A \Big| \mathbf{N}\mathbf{v} = \mathbf{0}, e_A + e_R + e_F + e_B + e_Q \le e_{tot}, \lambda = \frac{v_B}{e_{tot}} \text{ fixed}, \\ \forall j: v_j = k_j e_j f_j, e_j \ge 0, \{k_j, f_j, e_Q, e_{tot}\} \text{ constant} \Big\}, \end{aligned}$$

also given in eq. (10), is the formulation of the core model in chemostat conditions, where the growth rate is set by the experimentalist via the dilution rate.

In the regime $0 < \lambda < \lambda_c$, only the carbon-efficient mode is used, so the steady-state equations (1) and (5) admit a unique solution in terms of the protein fractions (eq. (17)). In the regime $\lambda_c < \lambda < \lambda_{max}$, a shift to the inefficient mode occurs because the $NG$-sector is depleted. Because the carbon assimilation rate is minimised, also the protein investment in the carbon-inefficient mode is minimised, since this mode requires more carbon due to its lower yield. So, the $NG$-sector remains empty across the regime $\lambda_c < \lambda < \lambda_{max}$. Again, this eliminates one of the variables and allows us to derive a unique solution to the steady-state equations in this regime, in terms of the protein fractions. We find that the corresponding expressions in terms of the model parameters are cumbersome. We use the expressions for the critical growth rate (19) and the maximal growth rate (22) to reformulate these as

$$\text{(S5)} \qquad \begin{aligned} \phi_B(\lambda) &= C_B\lambda \\ \phi_R(\lambda) &= C_R\left(\frac{\lambda_{max} - \lambda}{\lambda_{max}/\lambda_c - 1}\right) \\ \phi_F(\lambda) &= C_F\left(\frac{\lambda - \lambda_c}{1 - \lambda_c/\lambda_{max}}\right) \\ \phi_A(\lambda) &= \frac{\lambda_c\lambda_{max}(C_A^R - C_A^F)}{\lambda_{max} - \lambda_c} + \lambda\left(\frac{\lambda_{max}C_A^F - \lambda_c C_A^R}{\lambda_{max} - \lambda_c}\right). \end{aligned}$$

Here, the constants $C_j$ correspond to the same parameter combinations as in eq. (17) and (21).

The biosynthetic protein fraction $\phi_B$ satisfies the same linear relation with the growth rate in both regimes. The respiratory fraction $\phi_R(\lambda)$ attains its maximal value at the critical growth rate and vanishes at the maximal growth rate. Conversely, $\phi_F(\lambda)$ vanishes at the critical growth rate and attains its maximal value at the maximal growth rate. These results are summarised in Figure 2A. They agree with experimental findings [1; 2; 3].

In the regime $\lambda_c < \lambda < \lambda_{max}$, the flux vector is given by a convex combination (8) of the two EFMs $\mathbf{v}_R$ and $\mathbf{v}_F$ with nonzero convex coefficients $\alpha_R$ and $\alpha_F$. Because $\alpha_R(\lambda_c) = 1$ and $\alpha_F(\lambda_{max}) = 1$, we may express the fluxes through the lumped $R-$ and $F-$ reactions as the product of the corresponding convex coefficients and the maximal flux values. This yields

$$\begin{aligned} v_R(\lambda) &= \alpha_R(\lambda) v_R(\lambda_c) \\ v_F(\lambda) &= \alpha_F(\lambda) v_F(\lambda_{max}). \end{aligned} \tag{S6}$$

Expressions for the convex coefficients then follow directly from eq. (S5) as

$$\begin{aligned} \alpha_R(\lambda) &= \frac{v_R(\lambda)}{v_R(\lambda_c)} = \frac{\phi_R(\lambda)}{\phi_R(\lambda_c)} = \frac{\lambda_{max} - \lambda}{\lambda_{max} - \lambda_c} \\ \alpha_F(\lambda) &= \frac{v_F(\lambda)}{v_F(\lambda_{max})} = \frac{\phi_F(\lambda)}{\phi_F(\lambda_{max})} = \frac{\lambda - \lambda_c}{\lambda_{max} - \lambda_c}. \end{aligned} \tag{S7}$$

Here, we used that flux ratios are equivalent to protein fraction ratios when saturations are constant. These expressions for the convex coefficients satisfy the required property that $\alpha_R(\lambda) + \alpha_F(\lambda) = 1$.

### S4 Proteome efficiency definitions

In Section 3.3, we computed the proteome efficiencies for both modes as the ATP synthesis rate per unit of total protein expended to sustain the ATP flux. Using the solutions for the protein fractions (eq. (17), (21)), these condense to

$$\begin{aligned} \epsilon_R &\equiv \frac{m_R v_R(\lambda)}{e_B(\lambda) + e_R(\lambda) + e_A^R(\lambda)} = \frac{m_B}{C_B + C_R + C_A^R} = \frac{m_B}{1 - \phi_Q} \lambda_c \\ \epsilon_F &\equiv \frac{m_F v_F(\lambda)}{e_B(\lambda) + e_F(\lambda) + e_A^F(\lambda)} = \frac{m_B}{C_B + C_F + C_A^F} = \frac{m_B}{1 - \phi_Q} \lambda_{max}, \end{aligned}$$

If we assume constant enzyme saturations and ATP requirement $m_B$, the proteome efficiencies are also constant. Furthermore, these expressions show that the proteome efficiencies are directly related to the maximal growth rates: a shift to the inefficient $F$-mode thus implies its higher (maximal) growth rate as well as its higher proteome efficiency. In fact, taking the ratio of the proteome efficiencies shows that

$$\epsilon_{ratio} \equiv \frac{\epsilon_F}{\epsilon_R} = \frac{\lambda_{max}}{\lambda_c} \equiv \lambda_{ratio}. \tag{S8}$$

In other words, the break-even analysis in Section 3.5 yields identical results in terms the proteome efficiencies.

The proteome efficiency of the carbon-inefficient $F$-mode exceeds that of the carbon-efficient $R$-mode if its protein costs are lower, i.e.,

$$C_R + C_A^R > C_F + C_A^F. \tag{S9}$$

The biosynthetic protein costs $C_B$ do not appear, because these are the same for both strategies. This idea allows for a different definition for the proteome efficiency. Instead of considering the ATP synthesis rate per total amount of protein, an alternative is to compute the ATP synthesis rate per unit of catabolic protein expended to sustain the ATP flux. This yields the catabolic proteome efficiencies

$$\begin{aligned} \epsilon_R^{cat} &\equiv \frac{m_R v_R(\lambda)}{e_R(\lambda) + e_A^R(\lambda)} = \frac{m_B}{C_R + C_A^R} \\ \epsilon_F^{cat} &\equiv \frac{m_F v_F(\lambda)}{e_F(\lambda) + e_A^F(\lambda)} = \frac{m_B}{C_F + C_A^F}. \end{aligned}$$

These expressions cannot be simplified in terms of the critical and maximal growth rate. Instead, taking their ratio gives the inequality

$$\epsilon_{ratio}^{cat} \equiv \frac{\epsilon_F^{cat}}{\epsilon_R^{cat}} = \frac{C_R + C_A^R}{C_F + C_A^F} > \frac{C_B + C_R + C_A^R}{C_B + C_F + C_A^F} = \frac{\lambda_{max}}{\lambda_c} \equiv \lambda_{ratio}. \tag{S10}$$

A similar inequality was derived by Basan et al. [4] in their model for overflow metabolism in *E. coli.*

There is no reason to prefer one of the above definitions over the other, as both give a measure of the ATP synthesis rate per unit of protein. Furthermore, the condition $\epsilon_F > \epsilon_R$ is equivalent to $\epsilon_F^{cat} > \epsilon_R^{cat}$, because both are satisfied only when eq. (S9) holds. Regardless of the definition used, the proteome efficiencies do not encode additional information about the metabolic shift that is not already contained in the critical and maximal growth rates.

The various definitions presented here quantify the recent discussion on the concept of proteome efficiency and on the cause of overflow metabolism in yeasts [5; 6]. This will be analysed further by computing the proteome efficiencies in the yeast model in Supplementary Section S16.

### S5 Break-even analysis

In Section 3.5 we reformulated the break-even condition $\lambda_{ratio} = 1$ in terms of the model parameters as

$$\frac{N_R}{m_R k_R f_R} = \frac{N_F}{m_F k_F f_F} + \frac{(m_R - m_F)N_A}{m_R m_F k_A f_A}. \tag{S11}$$

Here, we use this relation to derive expressions for the curves depicted in the phase diagrams in Figure 3, that represent the break-even condition in terms of different parameter pairs.

Solving eq. (S11) for the ATP yield of the $R$-mode gives

$$m_R = \frac{m_F k_F f_F (k_R f_R + k_A f_A N_R)}{k_R f_R (k_F f_F + k_A f_A N_F)}. \tag{S12}$$

This equation represents the 'break-even point' for the ATP yield of the $R$-mode. It decreases with the catalytic constant of the $R$-mode ($k_R$), as also illustrated by Figure 3B. This means that either one or both these parameters should be small (compared to their counterpart for the $F$-mode) in order to observe the metabolic shift. On the other hand, the break-even point for $m_R$ increases linearly with the number of enzymatic reactions $N_R$ with some offset, as depicted in Figure 3C. This is because the protein costs $C_R$ increase with the number of reactions. So, a shift will occur only if the ATP generated per reaction by the carbon-efficient mode is small enough. Similar linear behaviour arises in Figure 3E when analyzing eq. (S12) as function of $m_F$. The break-even point for $m_R$ decreases with the number of reactions $N_F$, as presented in Figure 3F. Because increasing length $N_F$ increases the protein costs $C_F$, this also requires a lower ATP yield for the $R$-mode for a shift to occur. This relation is useful, as fermentation pathways may differ in length in various organisms. For example, *Clostridium pasteurianum* can ferment glucose to lactate or butyrate [7]. The latter involves six enzymatic reactions more.

We also solve eq. (S11) for the catalytic constant of the $R$-mode, giving its break-even point

$$k_R = \frac{m_F k_F f_F k_A f_A N_R}{f_R f_F k_F (m_R - m_F) + k_A f_A m_R f_R N_F)}. \tag{S13}$$

The break-even point for $k_R$ increases hyperbolically with the catalytic constant of the $F$-mode, as depicted in Figure 3D. This may be interpreted similar to the break-even relation between the ATP yields: when the catalytic constant of the $R$-mode increases, also that of the $F$-mode should increase to preserve the shift.

We can also use eq. (S11) to study relations between other parameter pairs that are not presented in Figure 3. For instance, eq. (S12) reveals that $m_R(k_F)$ is a hyperbolically increasing function, as presented in Figure S1A. Eq. (S13) and Figure S1B show that $k_R$ increases with $m_F$. For both relations, the interpretation is the same as for the catalytic constants in Figure 3D.

Instead of studying the break-even condition for parameters of the carbon-efficient $R$-mode, we can also inspect the parameters of the $F$-mode. Solving eq. (S11) for the ATP yield of the $F$-mode gives

$$m_F = \frac{m_R k_R f_R (k_F f_F + k_A f_A N_F)}{k_F f_F (k_R f_R + k_A f_A N_R)}. \tag{S14}$$

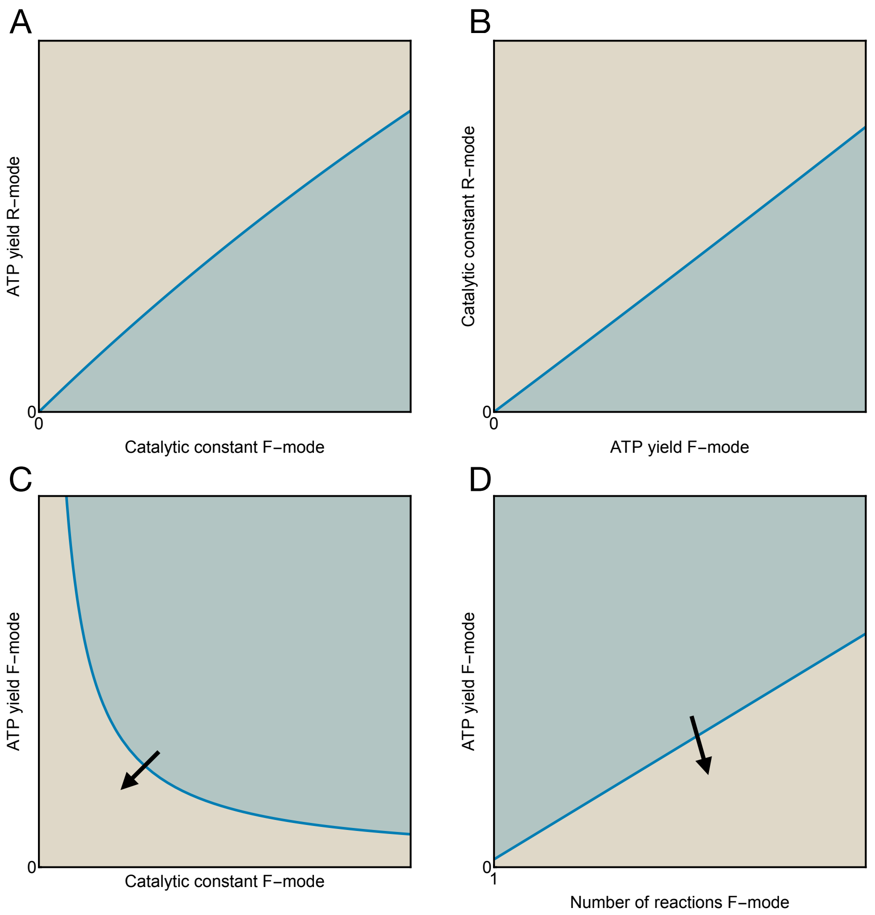


FIGURE S1. **Parameter perturbations in the model affect the occurrence of overflow metabolism.** The break-even analysis in terms of other parameter pairs that are not discussed in Section 3.5. For parameter configurations in the blue region, a metabolic shift occurs. In the brown region a shift will never occur. The arrow in C) and D) indicates the direction in which the break-even condition moves if the protein costs of the carbon-efficient mode increase.

This break-even point decreases as function of the catalytic constant $k_F$, while it increases as function of the number of reactions $N_F$. So, this gives the same behaviour as for the ATP-yield of the $R$-mode, but then with the shifting and non-shifting regions swapped. These two cases are presented in Figure S1C and D. The interpretation of these figures is also reversed with respect to Figure 3A and B, as increasing the protein costs $C_F$ will eliminate the shift. The arrow points in the direction that the regions move if the protein costs of the $R$-mode increase.

### S6 Analysis of metabolic shifts as function of nutrient quality during carbon uptake-limited growth

Here, we further analyse the case of uptake-limited growth on varying carbon sources for a constant titrated level of the uptake system. The quality of the different carbon sources may depend on various factors. First, we show that the catalytic rate constant of the assimilation pathway ($k_A$) is the best measure for the nutrient quality in our model. The concept of nutrient quality was introduced in Scott et al. [8] and recently studied by Mukherjee et al. [9]. Both studies state that the growth rate is proportional to the fraction of catabolic enzyme that catalyses the nutrient influx, in our model given by assimilation fraction

($\phi_A$), with the nutrient quality as proportionality constant. Studying the solutions for the assimilation fraction for both modes in eq. (17) and (21) shows that this proportionality constant is given by $C_A^{-1}$, the inverse of the protein costs for assimilation. The only parameters contained in the protein costs that may differ per carbon source are the catalytic rate constant ($k_A$) and the carbon required per mole of biomass synthesized ($Y_{X/B}$). The carbon requirement affects only the biosynthetic part of the metabolic network, and therefore does not affect the shift in the catabolic mode. This leaves $k_A$ as the only parameter that varies with carbon source that may affect the shift. This parameter thus represents the nutrient quality in our model.

We note that the concentration of the carbon source may also differ in uptake-limited conditions. This directly sets the saturation factor for assimilation ($f_A$). A more complete representation of the nutrient quality is therefore the product $k_A f_A$. For simplicity, we assume that the assimilation saturation factor remains fixed across different carbon sources, so we only vary $k_A$. Relaxing this assumption does not alter the analysis below.

### S6.1 Analytical solution to the optimisation problem in carbon uptake-limited conditions as function of the nutrient quality

The optimisation problem corresponding to this case of uptake-limited growth follows from adjusting eq. (9) by setting $e_A$ constant and $k_A$ as independent parameter, giving

(S15)
$$\max_{\mathbf{e}} \Big\{ \lambda = \frac{v_B}{e_{tot}} \Big| \mathbf{N}\mathbf{v} = \mathbf{0}, e_A + e_R + e_F + e_B + e_Q \le e_{tot}, k_A \text{ fixed}, \\ \forall j : v_j = k_j e_j f_j, e_j \ge 0, \{k_j, f_j, e_A, e_Q, e_{tot}\} \text{ constant} \Big\}.$$

This adjustment breaks the equivalence with the optimisation problem for chemostat conditions (eq. (10)), and thus requires a separate solution parameterized by $k_A$, that we will derive here.

Just as the other optimisations, eq. (S15) also maximises the biomass yield. At low nutrient quality, the carbon uptake rate and thus also the growth rate are low. In this regime, enzyme can be invested in growth-unassociated processes, hence the (equality) constraint on $e_{tot}$ in eq. (S15) is not active. As before, only the carbon-efficient mode will be used in this regime. We therefore study this pathway in isolation again, which yields a unique solution to the steady-state equations (1) and (5) in terms of the protein concentrations and the growth rate

(S16)
$$\begin{aligned} \lambda_R(k_A) &= Y^R_{B/S} f_A \frac{e_A}{e_{tot}} k_A \\ e^R_B(k_A) &= C_B Y^R_{B/S} f_A e_A k_A \\ e_R(k_A) &= C_R Y^R_{B/S} f_A e_A k_A \\ e_F(k_A) &= 0 \\ e_{NG}(k_A) &= e_{tot} - e_Q - e_A \left(1 + (C_B + C_R) Y^R_{B/S} f_A k_A\right). \end{aligned}$$

Here, we use the parameter combinations $C_j$ and the biomass yield from eq. (12) for notational convenience.

This solution is similar to the solution to eq. (10) in the regime $0 < \lambda < \lambda_c$ in eq. (17). Here, the growth-associated protein concentrations increase linearly with the nutrient quality instead of the growth rate. This diminishes the growth-unassociated pool $e_{NG}$, until it is depleted at a critical nutrient quality. This has the same interpretation as the critical growth rate. Solving $e_{NG} = 0$ for $k_A$ gives

(S17)
$$k^c_A = \frac{e_{tot} - e_Q - e_A}{(C_B + C_R) Y^R_{B/S} f_A e_A}.$$

The maximal nutrient quality $k^{max}_A$ follows from studying the carbon-inefficient mode in isolation, analogous to the derivation in Supplementary Section S2. We do not give the explicit solution in terms of the protein concentrations, as this follows directly from replacing all super- and subscripts $R$ in eq. (S16) by $F$. We find

(S18)
$$k^{max}_A = \frac{e_{tot} - e_Q - e_A}{(C_B + C_F) Y^F_{B/S} f_A e_A}.$$

For growth on a carbon source with maximal quality, the cell thus uses solely the carbon-inefficient mode. Furthermore, $k_A^{max} > k_A^c$ because $C_R > C_F$ and $Y_{B/S}^R > Y_{B/S}^F$. This is equivalent to the condition $\lambda_{max} > \lambda_c$, but reformulated in terms of the nutrient quality.

In the regime $k_A^c < k_A < k_A^{max}$, a convex combination of both modes is used. In this regime, $e_{NG} = 0$, so the optimisation problem (S15) again has an analytical solution for the protein concentrations. Here, we focus on the protein concentrations for the catabolic processes, as this is where the shift occurs. In the regime $k_A^c < k_A < k_A^{max}$, these are given by

$$\begin{aligned} e_R(k_A) &= C_R Y_{B/S}^R f_A e_A \left( \frac{k_A^{max} - k_A}{k_A^{max}/k_A^c - 1} \right) \\ e_F(k_A) &= C_F Y_{B/S}^F f_A e_A \left( \frac{k_A - k_A^c}{1 - k_A^c/k_A^{max}} \right). \end{aligned} \tag{S19}$$

These expressions are similar to their solution (S5) in chemostat conditions. The protein concentration $e_R$ decreases, while $e_F$ increases. This represents the gradual shift from the efficient to the inefficient mode as the nutrient quality increases. The corresponding fluxes are

$$\begin{aligned} v_R(k_A) &= k_R f_R e_R(k_A) = \frac{m_B}{m_R} Y_{B/S}^R f_A e_A \left( \frac{k_A^{max} - k_A}{k_A^{max}/k_A^c - 1} \right) \\ v_F(k_A) &= k_F f_F e_F(k_A) = \frac{m_B}{m_F} Y_{B/S}^F f_A e_A \left( \frac{k_A - k_A^c}{1 - k_A^c/k_A^{max}} \right). \end{aligned} \tag{S20}$$

We obtain the corresponding convex coefficients analogous to eq. (S7), giving

$$\begin{aligned} \alpha_R(k_A) &= \frac{v_R(k_A)}{v_R(k_A^c)} = \frac{e_R(k_A)}{e_R(k_A^c)} = \frac{k_A^{max} - k_A}{k_A^{max} - k_A^c} \\ \alpha_F(k_A) &= \frac{v_F(k_A)}{v_F(k_A^{max})} = \frac{e_F(k_A)}{e_F(k_A^{max})} = \frac{k_A - k_A^c}{k_A^{max} - k_A^c}. \end{aligned} \tag{S21}$$

Similarly, these expressions satisfy $\alpha_R(k_A) + \alpha_F(k_A) = 1$.

#### S6.2 Analytical solution to the optimisation problem in carbon uptake-limited conditions as function of the growth rate

The catalytic constant of the assimilation pathway may be difficult to obtain experimentally. Also, the critical nutrient quality (S17) and maximal nutrient quality (S18) depend significantly on the constant titrated concentration of the assimilation enzyme. Instead, we may express the solution to eq. (S15) in the regime $k_A^c < k_A < k_A^{max}$ in terms of quantities that are easier to obtain experimentally, such as the growth rate $\lambda$.

The corresponding critical growth rate is obtained from $\lambda_R(k_A)$ in eq. (S16) by substituting the critical nutrient quality $k_A^c$. Similarly, the corresponding maximal growth rate follows from substituting the maximal nutrient quality $k_A^{max}$ in $\lambda_F(k_A)$ (not shown here), which is the solution for the growth rate when the inefficient mode is used in isolation. This yields

$$\begin{aligned} \lambda_R(k_A^c) &= \frac{1 - \phi_Q - \phi_A}{C_B + C_R} \\ \lambda_F(k_A^{max}) &= \frac{1 - \phi_Q - \phi_A}{C_B + C_F}. \end{aligned} \tag{S22}$$

These expressions differ from the critical growth rate (19) and the maximal growth rate (22) obtained for chemostat conditions, because the constant assimilation fraction $\phi_A$ now appears in the numerator. Reformulating the protein concentrations in eq. (S19) in terms of the growth rate gives

$$\begin{aligned} e_R(\lambda) &= C_R e_{tot} \left( \frac{\lambda_F(k_A^{max}) - \lambda}{\lambda_F(k_A^{max})/\lambda_R(k_A^c) - 1} \right) \\ e_F(\lambda) &= C_F e_{tot} \left( \frac{\lambda - \lambda_R(k_A^c)}{1 - \lambda_R(k_A^c)/\lambda_F(k_A^{max})} \right). \end{aligned} \tag{S23}$$

These protein concentrations only differ from their analogous solutions (S5) in chemostat conditions through the different expressions for the critical and maximal growth rate in eq. (S22). So, the general mathematical structure of the solution to the optimisation problems remains conserved across different experimental conditions.

The overflow product flux is proportional to the flux through the $F$ process ($v_F$). Reformulating the solution for $v_F$ in eq. (S20) as function of the growth rate gives

$$v_F(\lambda) = k_F f_F e_F(\lambda) = \frac{m_B}{m_F} e_{tot} \left( \frac{\lambda - \lambda_R(k_A^c)}{1 - \lambda_R(k_A^c)/\lambda_F(k_A^{max})} \right). \tag{S24}$$

This shows that the overflow flux is a linear function of the growth rate, both for chemostat and carbon uptake-limited batch conditions. Furthermore, both its slope and intersection with the $\lambda$-axis are directly determined by the critical and the maximal growth rates. In the limit of small $\phi_A$, the expressions for these growth rates in eq. (S22) reduce to the analogous expressions in chemostat conditions, respectively in eq. (19) and (22), for low assimilation costs ($C_A$). In other words, the overflow line appearing in both chemostat cultures and in uptake-limited batch cultures in different nutrient conditions can be explained using the same principles.

### S7 Construction of the core model for yeast

We adapt the core model to *S. cerevisiae* cultivated in aerobic glucose-limited chemostats. The corresponding reaction network is depicted in Figure 4. Here, the uptake of carbon is described by a separate transport reaction $T$. Glycolysis $G$ degrades glucose into pyruvate, thereby gaining $m_G$ ATP per glucose. Pyruvate is then degraded into byproducts by one of the two catabolic modes. Respiration $R$ degrades pyruvate into carbon dioxide and water, which yields an additional $m_R$ ATP per glucose via the TCA cycle and oxidative phosphorylation. Ethanol fermentation $F$ degrades pyruvate into ethanol. The concentrations for the internal metabolites (ATP, glucose and pyruvate) satisfy the differential equations

$$\begin{aligned} \dot{ATP} &= m_G v_G + m_R v_R - m_B v_B \\ \dot{Glc}_{in} &= v_T - v_G \\ \dot{Pyr} &= v_G - v_F - v_R. \end{aligned} \tag{S25}$$

The equation for pyruvate does not include a pyruvate flux towards biosynthesis, because we assume that biosynthesis does not require carbon. A different approach is to assume a constant biosynthetic carbon demand, as in the general core model. However, the biosynthetic carbon demand may change when the cell shifts from respiration to fermentation. Accounting for this effect is beyond the scope of this work, because we assume that catabolism and anabolism are independent. We therefore decided to neglect biosynthetic carbon entirely, and adjust for this after evaluating the model. This is discussed in Supplementary Section S11.

The differential equations in eq. (S25) may be reformulated in terms of a flux vector $\mathbf{v_y} = (v_T\ v_G\ v_R\ v_F\ v_B)^T$ and the corresponding stoichiometric matrix

$$\mathbf{N_y} = \begin{pmatrix} 0 & m_G & m_R & 0 & -m_B \\ 1 & -1 & 0 & 0 & 0 \\ 0 & 1 & -1 & -1 & 0 \end{pmatrix}. \tag{S26}$$

We assume balanced growth conditions, where metabolism operates at steady state, i.e., $\mathbf{N_y v_y} = \mathbf{0}$. The two EFMs of the reaction network in Figure 4 correspond to respiration and fermentation. They follow from the stoichiometric matrix as

$$\mathbf{v}_{res} = v_B \begin{pmatrix} \frac{m_B}{m_R+m_G} \\ \frac{m_B}{m_R+m_G} \\ \frac{m_B}{m_R+m_G} \\ 0 \\ 1 \end{pmatrix}, \quad \mathbf{v}_{fer} = v_B \begin{pmatrix} \frac{m_B}{m_G} \\ \frac{m_B}{m_G} \\ 0 \\ \frac{m_B}{m_G} \\ 1 \end{pmatrix}. \tag{S27}$$

Both EFMs have a biomass yield that we compute by taking the ratio of the last over the first entry (following eq. (11)), giving

$$Y_{cat}^{res} = \frac{m_R + m_G}{m_B}, \quad Y_{cat}^{fer} = \frac{m_G}{m_B}. \tag{S28}$$

Because carbon for biosynthesis is not included, these yields describe the biomass yield per mole of glucose that is used by catabolism. The higher biomass yield of respiration directly follows from its higher ATP yield.

The total protein concentration in a yeast cell is assumed fixed [10] and decomposed into the protein sectors as

$$e_{tot} = e_T + e_G + e_R + e_F + e_B + e_{NG} + e_Q. \tag{S29}$$

Next to $e_{tot}$, we also assume that the growth-unassociated $Q$-sector and the transporter enzyme have a fixed concentration [3; 8].

We implement the yeast model as an optimisation problem by minimising the carbon transport rate for variable protein concentrations at a fixed growth rate. Because the transporter enzyme concentration is assumed fixed, minimisation of the carbon transport rate reduces to minimisation of the transporter saturation factor. The optimisation problem for the yeast model is then given by

$$\begin{aligned} \min_{\mathbf{e}} \Big\{ f_T \big| \mathbf{N_y v_y} = \mathbf{0}, e_T + e_G + e_R + e_F + e_B + e_{NG} + e_Q = e_{tot}, \lambda = \frac{v_B}{e_{tot}} \text{ fixed}, f_T \geq 0, \\ \forall j : v_j = \langle k \rangle_j \frac{e_j}{N_j} \langle f(\lambda) \rangle_j, e_j \geq 0 \text{ and } \{\langle k \rangle_j, N_j, e_T, e_Q, e_{tot}\} \text{ constant} \Big\}. \end{aligned} \tag{S30}$$

The equations for the lumped reaction rates $v_j$ are derived in the next section. Model parameters and their values are presented in Table S6. A Mathematica implementation of the yeast model is provided in the Supporting Information.

### S8 Protocol for reducing GEMs to core models

The core model presented in this work is not tailored to any metabolic shift in a specific organism. Adapting this model to a specific organism requires identification of the essential metabolic pathways and protein sectors that play a significant role during the shift. Each growth-associated sector $j$ is then described by a lumped reaction with flux $v_j$. These fluxes may be expressed in terms of kinetic parameters of the individual enzymatic reactions that are contained in the sector. These parameters can be obtained from a GEM integrated with proteomics data, thereby calibrating the model with experimental data. Using this top-down approach, we reduce the large metabolic network from the GEM to a core metabolic network, such as the one presented here. Erdrich et al. [11] use another method for automated reduction of GEMs. Their method, however, typically results in a core model that exceeds manageable size for the mathematical analysis conducted in this study.

In this section, we derive expressions for the fluxes $v_j$ in the core model adapted to yeast. This method is straightforward to generalize to other unicellular organisms. We start from the complete metabolic network of yeast that consists of $N$ enzyme-catalysed reactions. The rate of reaction $i$ is given by $v_i = k_i e_i f_i$. We assume that each enzyme $e_i$ belongs to only one of the seven protein sectors. In other words, we may decompose the set of $N$ metabolic reactions as disjoint subsets per sector. Each subset consists of the $N_j$ individual enzymatic reactions that are contained in sector $j$, such that $N = \sum_j N_j$. The total protein concentration then decomposes as

$$e_{tot} = \sum_{i=1}^{N} e_i = e_T + \sum_{i=1}^{N_G} e_i^G + \sum_{i=1}^{N_R} e_i^R + \sum_{i=1}^{N_F} e_i^F + \sum_{i=1}^{N_B} e_i^B + \sum_{i=1}^{N_{NG}} e_i^{NG} + \sum_{i=1}^{N_Q} e_i^Q. \tag{S31}$$

We assume that there is only one transport reaction. We then define the enzyme concentrations per sector as

$$e_j \equiv \sum_{i=1}^{N_j} e_i^j, \tag{S32}$$

such that we retrieve eq. (S29).

Now, we focus on the growth-associated protein sectors, since we describe only these by a lumped reaction with flux $v_j$. Due to variations in reaction stoichiometry, flux ratios within sectors are not necessarily equal to one. For instance, each time the upper part of glycolysis runs, the lower part has to run twice to preserve steady-state. These flux ratios within sectors we call the *intrasector* stoichiometries. These are different from the *intersector* stoichiometries, which are defined by the entries of the EFMs in eq. (S27).

In principle, the intrasector stoichiometries may be obtained per sector. This is straightforward for linear metabolic pathways such as glycolysis, but it becomes more involved when considering sectors that consist of more complex pathways, such as respiration or biomass synthesis. Furthermore, these intrasector stoichiometries may differ per EFM. Obtaining these stoichiometries is difficult when a combination of EFMs is used, which is the main regime of interest in this study. Therefore, we disregard the intrasector stoichiometries by assuming that all fluxes within a sector are identical, i.e., $v_i^j = v_j$ for all reactions in sector $j$. Rewriting eq. (S32) by using the rates $v_i = k_i e_i f_i$ and applying this assumption gives

$$e_j = \sum_{i=1}^{N_j} e_i^j = \sum_{i=1}^{N_j} \frac{v_i^j}{k_i^j f_i^j} = v_j \sum_{i=1}^{N_j} \frac{1}{k_i^j f_i^j} = v_j N_j \left\langle \frac{1}{kf} \right\rangle_j . \tag{S33}$$

Here, $\langle ... \rangle_j$ denotes the arithmetic mean over the reactions in sector $j$. Reformulating this gives an equation for the lumped reaction fluxes

$$v_j = \left\langle \frac{1}{kf} \right\rangle_j^{-1} \frac{e_j}{N_j}. \tag{S34}$$

The ratio $\frac{e_j}{N_j}$ represents the average enzyme concentration per reaction in sector $j$.

Because we assume that carbon transport consists of only one reaction, its flux trivially reduces to

$$v_T = \left( \frac{1}{k_T f_T} \right)^{-1} e_T = k_T e_T f_T. \tag{S35}$$

For the other growth-associated sectors $j \in \{G, R, F, B\}$ that contain more reactions we derive alternative approximation for the lumped reaction fluxes $v_j$. First, we assume that the catalytic rate constants $k_i$ and saturations $f_i$ are independent parameters. This means that these parameters do not covary. This results in the approximation

$$v_j^H = \left\langle \frac{1}{k} \right\rangle_j^{-1} \frac{e_j}{N_j} \left\langle \frac{1}{f} \right\rangle_j^{-1} \tag{S36}$$

that expresses the lumped reaction fluxes in terms of the harmonic means of $k_i$ and $f_i$. Second, we can use the general relation that arithmetic means are always larger than harmonic means. In other words, we have a lower bound $\langle k \rangle_j \geq \left\langle \frac{1}{k} \right\rangle_j^{-1}$, $\langle f \rangle_j \geq \left\langle \frac{1}{f} \right\rangle_j^{-1}$ for the arithmetic means. This gives

$$v_j^A = \langle k \rangle_j \frac{e_j}{N_j} \langle f \rangle_j \geq v_j^H. \tag{S37}$$

This expresses the flux through lumped reaction $j$ as a product of the mean catalytic rate constant, the mean saturation factor and the mean enzyme concentration per reaction in the sector. Therefore, $v_j^A$ should be interpreted as the mean flux through sector $j$.

The question remains which of these three approximations (S34), (S36) or (S37), is the best to use for the core model, i.e., which of these models the (mean) flux through the sectors most accurately. The first approximation (S34) is not convenient, because this makes use of products of catalytic constants and saturation factors for individual reactions. Catalytic constants and protein concentrations (from which saturations can be computed, as explained in the following section) are, however, typically measured as separate quantities during experiments. Representing them separately allows us to use their individual properties, such as the fact that saturation factors can not exceed one, when calibrating the model with experimental data.

Because of the inequality in eq. (S37), the approximation in terms of harmonic means in eq. (S36) appears to be more accurate. However, the accuracy of these approximations also depends on the intrasector stoichiometries, which are not taken into account. If the average intrasector stoichiometric coefficient exceeds one, the approximation in terms of arithmetic means might be more accurate than the approximation in terms of harmonic means. So, *a priori*, there is no justification to prefer one of these approximations for the lumped reaction fluxes.

Therefore, we should establish a different criterion to select the best approximation for the lumped reaction fluxes in the yeast model. Two key characteristics of overflowing yeast are the critical dilution rate $D_c = 0.275\,\mathrm{h}^{-1}$ in glucose-limited chemostats and the maximal growth rate in glucose batch, which is $\lambda_{max} = 0.47\,\mathrm{h}^{-1}$ for the strain used in Elsemman et al. [3]. The yeast model should be calibrated to these two growth rates. Following the methods of the next section, we computed the harmonic means and arithmetic means of the catalytic rate constants and the enzyme saturations and ran the model in Mathematica for both approximations. Using the arithmetic means results in a critical and maximal growth rate that are in the right order of magnitude, while using the harmonic means is less accurate. Based on this analysis, we decided to use the approximation of the lumped reaction fluxes in eq. (S37) in terms of the arithmetic means. Further calibration of model parameters to find the observed critical and maximal growth rate is discussed in Supplementary Section S10.

### S9 Saturation functions and catalytic constants in the yeast model

The remaining parameters of the yeast model are determined from proteomics data from Elsemman et al. [3]. These are the mean catalytic rate constants $\langle k \rangle_j$, the mean enzyme saturations $\langle f \rangle_j$ and the number of individual enzymatic reactions $N_j$ per sector.

#### S9.1 Pathway categorisation and number of reactions per sector

First, the metabolic pathways and corresponding enzymatic reactions included in the GEM from Elsemman et al. [3] need to be categorised in terms of the protein sectors used in the yeast model. This categorisation, given in Table S1, is based on the COG categories [12]. We only include enzymatic reactions that are given a reaction ID, as these are the reactions that are considered to be metabolic. This results in a set of reactions $I_j$ that contains all enzymatic reactions from the GEM that are associated with the (coarse-grained) sector $j$. For example, the biomass synthesis sector contains all pathways and corresponding reactions that are related to protein synthesis, because we assume that biomass consists only of proteins.

Intuitively, the number of enzymatic reactions $N_j$ per sector $j$ should correspond to $|I_j|$, the number of enzymatic reactions in each set $I_j$. However, these reaction sets also contain other involved reactions that do not directly contribute to the net molecular conversion performed by the sector. The core model only describes metabolic processes that are essential to model the metabolic shift. Therefore, it should only take distinct molecular conversions into account. The number of reactions $|I_j|$ in the reaction sets is not an accurate representation of the distinct molecular conversions that should be modeled for these sectors. Instead, these lengths $N_j$ are obtained from standard literature and given in Table S2. The number of reactions required for biomass synthesis exceeds that of all other sectors, but it is not exactly known. Therefore, we do not include a value in Table S2. Instead, we use this parameter to calibrate the model to experimental data, as is discussed further in Supplementary Section S10.

| **Sector in the yeast model** | **Included pathways from [3]** |
|---|---|
| Transport | Glucose transport |
| Glycolysis | Glycolysis |
| Respiration | Mitochondrial carriers, Mitochondrial ribosome, Oxidative phosphorylation, TCA cycle, TIM/TOM translocase |
| Fermentation | Ethanol fermentation |
| Biomass synthesis | Amino acid biosynthesis, Amino acid-tRNA ligases, Protein folding, Ribosome, Ribosome assembly factors, Translation elongation factors, Translation initiation factors, tRNA modification |

TABLE S1. **Metabolic pathways from the GEM in Elsemman et al. [3], categorised per protein sector in the yeast model.** Biomass synthesis contains all pathways related to protein synthesis, of which the ribosome is a subset.

Although not all reactions in the sets $I_j$ are taken into account to compute the number of distinct conversions $N_j$, all these reactions do contribute to the mean kinetic constant and mean enzyme saturation of the sectors. All these reactions are therefore included to compute these means. An exception to this is fermentation, for which we only include the two canonical fermentation reactions that are catalysed by the enzymes pyruvate decarboxylase and alcohol dehydrogenase to compute the mean parameters. This manual curation for fermentation was possible, since $|I_F| = 7$. In principle, such a manual check should be carried out for all sectors. However, this is infeasible and will likely not give significantly different results for the other sectors.

| Parameter symbol | Included pathways | Number of reactions |
|---|---|---|
| $N_T$ | Glucose transport | 1 |
| $N_G$ | Glycolysis | 10 |
| $N_R$ | Oxidative phosphorylation, TCA cycle, Mitochondrial carriers | 20 |
| $N_F$ | Ethanol fermentation | 2 |

Table S2. **Table with number of reactions per process in the yeast model.** Because $N_T = 1$, this parameter is not used throughout the rest of this work. In yeast, oxidative phosphorylation is separated into five reaction steps. The TCA cycle consists of the eight standard reactions. The number of reactions related to mitochondrial carriers (7) is obtained from Elsemman et al. [3].

### S9.2 Computing saturations for individual reactions and special cases

Changes in environmental conditions affect intracellular concentrations and thus also enzyme saturations. Some studies explicitly model enzyme kinetics and reactant concentrations to account for changing saturations [13; 14]. However, for coarse-grained models, this usually shows only qualitative agreement between the model and experimental data. A different approach is to infer enzyme saturations from proteomics data. Elsemman et al. [3] report measured protein mass fractions $e_i^{exp}$ from chemostat experiments and GEM-predicted protein mass fractions $e_i^{GEM}$. We convert these to concentration fractions by dividing the mass fractions by the molar protein mass. The enzyme saturations now follow from assuming that the GEM predicts the right fluxes, i.e., we assume that $v_i^{GEM} = v_i^{exp}$, where $v_i^{exp}$ is the actual flux through reaction $i$. The GEM predicts a minimal enzyme investment ($e_i^{GEM}$) required to sustain this flux, as it sets all enzyme saturations to one. If we additionally assume that the GEM contains the correct catalytic constants, i.e., $k_i^{GEM} = k_i^{exp}$, then the enzyme saturation of reaction $i$ is

$$f_i^{exp} = \frac{e_i^{GEM}}{e_i^{exp}}. \qquad \text{(S38)}$$

As emphasised earlier, the number of reactions $|I_j|$ in the reaction set of process $j$ usually does not correspond to the number of distinct molecular conversions $N_j$. This is partially due to conversions that can not be described by a single reaction catalysed by one enzyme. Such special conversions also require a slightly different computation of the corresponding saturation. In the yeast model, we account for two of these special cases.

The first case concerns multimeric enzymes, which are protein complexes that consist of multiple subunits. Although the complex catalyses the reaction, the subunits appear as separate proteins in the proteomics data. Since the data do not distinguish between free or bound subunits, we assume that all subunits are bound into the catalysing complex. If a subunit with concentration $e_{im}$ appears multiple times in a complex with concentration $e_i^{complex}$, it is assigned a stoichiometric coefficient $n_{im}$. The subunit with the lowest concentration (after normalising for the stoichiometries) limits formation of the complex. So, the minimum of the individual subunit concentrations gives an approximation for the concentration of the complex

$$e_i^{complex} \approx \min_m \frac{e_{im}}{n_{im}}. \qquad \text{(S39)}$$

For each multimeric enzyme, this approximation is used to compute its concentration from the data for the individual subunits. This calculation is done for both the experimental data and the GEM predictions from Elsemman et al. [3]. Plugging in these concentrations in eq. (S38) then gives the (approximated) saturation of each multimeric enzyme.

Second, enzymes exist that catalyse the same reaction. These are called iso-enzymes. Because enzyme saturations are determined by the concentrations of substrate and products of the reaction, these iso-enzymes should have the same saturation factor. For reactions catalysed by multiple iso-enzymes, the total concentration of catalysing enzymes ($e_i^{iso}$) is obtained by summing the concentrations $e_{in}$ of all iso-enzymes $n$ as

$$e_i^{iso} = \sum_n e_{in}. \tag{S40}$$

Again, this is computed for both the experimental data and for the GEM predictions from [3]. Plugging in these concentrations in eq. (S38) gives the saturation of the reaction catalysed by iso-enzymes. Some reactions are catalysed by multiple iso-enzymes that are also multimeric. In that case, both cases need to be accounted for.

### S9.3 Determination of saturation functions

The proteomics data in Elsemman et al. [3] is obtained from glucose-limited chemostat experiments set at different dilution rates, which we call $D_l$. The GEM was run at slightly different dilution rates $D_k$ by the authors. For each $D_l$, we find the $k$ such that $D_k$ is closest to $D_l$. Thereby we match the experimentally studied dilution rates to the simulated ones. Using eq. (S38), this results in an array $f_{il} = f_i(D_l)$ for the saturation of each reaction $i$ at dilution rates $D_l$. For some reactions, experimental data shows expression of the catalysing enzyme, while the GEM predicted its concentration to be zero. These reactions are neglected because their saturations can not be computed from eq. (S38). In other cases, the computed saturation exceeds one. By definition, this is no feasible value for the saturation factor. These cases are likely caused by an underestimation of the catalytic constant $k_i^{GEM}$. This results in a predicted concentration $e_i^{GEM}$ that exceeds the measured $e_i^{exp}$, even at maximal predicted enzyme saturation. We adjust these underestimated catalytic constants by increasing them until $f_{il} \leq 1$ for all dilution rate $D_l$. This results in both feasible saturation values and (more) realistic estimations of the effective catalytic constants. A similar adjustment is used by Battjes et al. [5].

The saturation arrays $f_{il}$ are then grouped per reaction set $I_j$ according to the categorisation in Table S1. For each sector $j$ in the yeast model, we then compute the arithmetic mean of the saturation per individual reaction at each dilution rate $D_l$. This results in arrays with elements

$$\langle f_l \rangle_j = \sum_{i \in I_j}^{|I_j|} \frac{f_{il}}{|I_j|}. \tag{S41}$$

The resulting dataset is provided in the Supporting Information. We plot this data as function of the growth rate (which equals the dilution rate in the chemostat) and fit functions $\langle f(\lambda) \rangle_j$ to it. These functions we refer to as the saturation functions.

The data $\langle f_l \rangle_j$ together with the fitted saturation functions are depicted in Figure S2. The mean saturation data for glycolysis show distinct linear behaviour before and after the critical dilution rate. Therefore, we have fit a piecewise linear function to this data. The saturation of enzymes used for ethanol fermentation is close to zero at low dilution rates, because ethanol is produced only after the critical dilution rate. For higher dilution rates, the saturation increases hyperbolically. This behaviour is also captured by a piecewise continuous function. Figure S2C shows that a clear trend is missing in the mean saturation of respiration. Obtaining a saturation function $\langle f(\lambda) \rangle_R$ from this data is problematic. Therefore, we assume that $\langle f(\lambda) \rangle_R$ is a hyperbolically increasing function. This results in a monotonic specific oxygen uptake rate (before and after the critical growth rate) that is consistent with experimental data [3]. A different approach is to use a piecewise continuous function that exhibits a drop around the critical dilution rate. This is a better fit to the saturation data. However, it gives results for the respiratory protein fraction and oxygen uptake rate that fit less well to experimental data. Therefore, we use the hyperbolically increasing form for $\langle f(\lambda) \rangle_R$ in

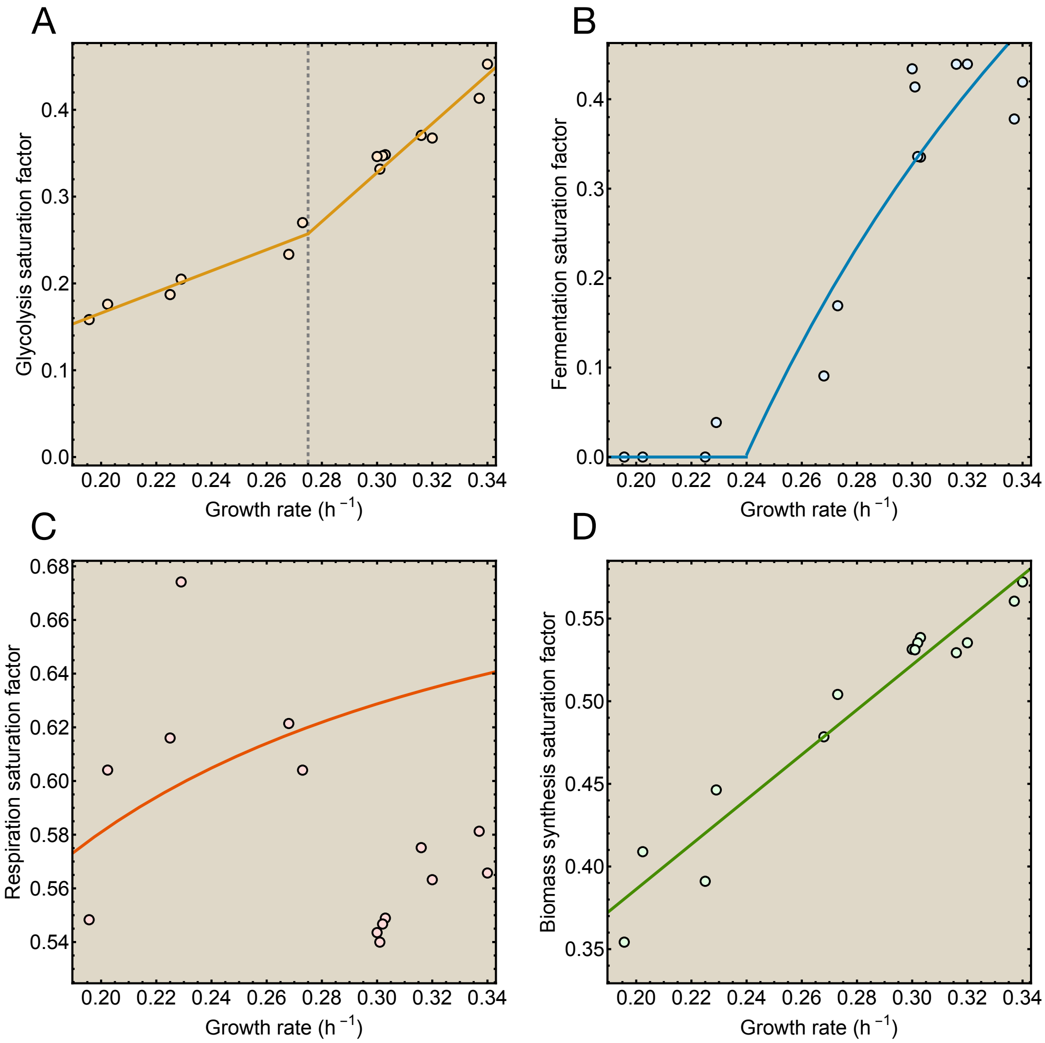


Figure S2. **Saturation functions used in the yeast model.** Saturation functions are fit to data obtained from proteomics data, adjusted with permission from [3] (Creative Commons Attribution 4.0 International License). The saturation functions describe the mean saturation factor of the four growth-associated processes in the yeast model. The saturation functions increase monotonically with the growth rate.

the rest of this work. For the enzymes involved in biomass synthesis, we obtain a linear saturation function $\langle f(\lambda)\rangle_B$ from the data.

#### S9.4 Catalytic rate constants

The GEM in Elsemman et al. [3] makes use of (estimated) catalytic rate constants $k_i^{GEM}$ to compute reaction fluxes. We obtain the mean catalytic rate constants $\langle k\rangle_j$ by computing the arithmetic mean of the catalytic constants per sector as

$$\langle k\rangle_j = \sum_{i\in I_j}^{|I_j|} \frac{k_i^{GEM}}{|I_j|}. \tag{S42}$$

Here, we include the effective $k_i^{GEM}$ that we get by increasing the underestimated catalytic constants so that all saturation values remain below one. We omit proteins for which no value of the catalytic constant is reported. The resulting mean catalytic constants are given in Table S3.

For biomass synthesis, we do not use the mean catalytic constant of the reactions in this sector as parameter in the yeast model. This is because we assume that biomass consists only of protein. This means that the biomass synthesis rate $v_B$ represents the protein synthesis rate. The corresponding catalytic constant is the protein translation rate by the ribosome, or more specifically its peptide elongation rate $k_{ribo}$.

| **Sector in the yeast model** | **Catalytic constant symbol** | **Catalytic constant value ($h^{-1}$)** |
|---|---|---|
| Transport | $\langle k \rangle_T$ | $7.2 \cdot 10^5$ |
| Glycolysis | $\langle k \rangle_G$ | $1.19 \cdot 10^6$ |
| Respiration | $\langle k \rangle_R$ | $4.35 \cdot 10^6$ |
| Fermentation | $\langle k \rangle_F$ | $1.68 \cdot 10^6$ |
| Biomass synthesis | $k_{ribo}$ | 126 |

Table S3. **Mean catalytic rate constant for each sector in the yeast model.** For $k_{ribo}$, we use the value reported by Elsemman et al. [3].

### S10 Calibration of the yeast model

Using the parameter values from the previous sections, we compute the solution to the optimisation problem (S30) for yeast numerically in Mathematica. The results show a shift from respiration to fermentation. However, the dilution rate when the shift initiates does not correspond to the observed critical dilution rate $D_c = \lambda_c^y = 0.275\,\mathrm{h}^{-1}$ for yeast in aerobic glucose-limited chemostats. Also, the computed maximal growth rate is inconsistent with the observed value $\lambda_{max}^y = 0.47\,\mathrm{h}^{-1}$ for the strain used in Elsemman et al. [3]. These discrepancies may be due to the size of the core model, or the approximation of fluxes in eq. (S37) in terms of arithmetic means of parameters (although this gave more accurate results than using the approximation in eq. (S36) in terms of harmonic means). Finding the observed values for the critical and maximal growth rate in the yeast model therefore requires additional parameter calibration.

Some of the catalytic constants of individual enzymatic reactions were underestimated in the GEM, as discussed in Supplementary Section S9. We increased these cases appropriately to the effective values. These effective values are more realistic, but this method does not guarantee that they are exactly correct. So, this suggests that the (mean) catalytic constants (S42) are parameters with high uncertainty. Therefore, we argue that these are the appropriate parameters to adjust in order to calibrate the model. The catalytic constant of biomass synthesis is set as the peptide elongation rate $k_{ribo}$ and is obtained from the literature, so this parameter is not adjusted. We also use the number of biosynthetic reactions $N_B$ as calibration parameter, since it is not possible to obtain its exact value.

The calibrated values of these parameters are given in Table S4. It should be noted that the catalytic constant of respiration requires a 100-fold decrease with respect to its mean value $\langle k \rangle_R$. This likely indicates that $\langle k \rangle_R$ is a poor representation of the mean catalytic constant of respiration. Another explanation may be that respiratory fluxes can not be approximated well in terms of mean enzyme-kinetic parameters.

| **Parameter description** | **Parameter symbol** | **Calibrated value** |
|---|---|---|
| Catalytic constant of glycolysis | $k_G^{cal}$ | $0.43 \langle k \rangle_G$ |
| Catalytic constant of respiration | $k_R^{cal}$ | $0.0077 \langle k \rangle_R$ |
| Catalytic constant of fermentation | $k_F^{cal}$ | $1.1 \langle k \rangle_F$ |
| Number of biosynthetic reactions | $N_B^{cal}$ | 142 |

Table S4. **Calibrated parameters and their values in the yeast model.**

### S11 Adjustment and results for carbon uptake in the yeast model

In the yeast model, we only account for glucose that is degraded into byproducts by the catabolic modes that charge ATP. However, glucose is also required to synthesize precursors for biomass. This leads to a model prediction for the glucose uptake flux that is significantly below the observed uptake. To compare the results of the yeast model with experimental data, we need to correct for this unaccounted biosynthetic carbon. We do this by adjusting the specific glucose uptake flux predicted by the model, represented by the dashed orange line in Figure S3A. We increase the predicted glucose uptake rate such that it matches the experimental data from Elsemman et al. [3] at the critical and maximal growth rate. The resulting adjustment factor is the fraction of imported glucose that is used for catabolic processes ($\frac{S_{cat}}{S_{tot}}$), depicted in Figure S3B. This fraction is constant before the critical growth rate, because the flux ratios (and thus yields) are constant when only one EFM is used. The catabolic glucose fraction increases in the regime $\lambda_c < \lambda < \lambda_{max}$. This follows from the lower ATP yield per mole glucose of fermentation. Multiplying the dashed orange line in Figure S3A by the inverse of the catabolic glucose fraction ($\frac{S_{tot}}{S_{cat}}$) results in the solid orange line. This represents the total specific glucose uptake flux, that agrees better with the experimental data.

We apply the same adjustment to the biomass yield. First, we convert the yield obtained from the model to units of gram dry weight per gram glucose. This conversion uses the molar weight of an average protein $MW_{prot} = 50$ g/mmol protein (since we assume biomass consists only of proteins) and the molar weight of glucose $MW_{Glc} = 0.180$ g/mmol glucose. The resulting 'catabolic' yield $Y^{cat}_{B/S}$ predicted by the model is shown by the dashed green line in Figure S3C. It exceeds the measured yield at all growth rates. To obtain a more realistic prediction for the yield, we multiply the catabolic yield by the catabolic glucose fraction, giving

$$Y^{cat}_{B/S}(\lambda)\frac{S_{cat}}{S_{tot}} = \frac{v_B}{v_T}\frac{S_{cat}}{S_{tot}} = \frac{B\,h^{-1}}{S_{cat}\,h^{-1}}\frac{S_{cat}}{S_{tot}} = \frac{B}{S_{tot}} = Y^{tot}_{B/S}(\lambda). \tag{S43}$$

The solid green line in Figure S3C represents the biomass yield per total amount of glucose consumed. This also agrees better with the experimental data.

### S12 Adjustment and results for oxygen uptake and ATP demand in the yeast model

Respiration requires six moles of oxygen for the complete oxidation of one mole of glucose to carbon dioxide and water. Since the respiratory flux $v_R$ has a unit of moles glucose per hour in the yeast model, the predicted specific oxygen uptake flux is $q_{O_2} = 6\frac{v_R}{\rho_{DW}}$. This is represented by the dashed red line in Figure S3D. The model underestimates the oxygen uptake rate compared to the experimental data for two reasons.

The first considers the carbon flow through respiration. We assume that all this carbon is converted into carbon dioxide. However, some reactions in the TCA cycle also produce precursors for biosynthesis, like $\alpha$-ketoglutarate. Taking this into account would increase the respiratory flux, and thus also increase the oxygen uptake rate. However, this would require an assessment of the carbon flow per individual reaction, which is outside the scope of this coarse-grained model.

The second factor that affects the oxygen uptake flux is the ATP demand. Since both glycolysis and the TCA cycle also provide precursors for biosynthesis (such as, respectively, pyruvate and $\alpha$-ketoglutarate), this makes these pathways in part biosynthetic. Therefore, a fraction of the ATP generated by these pathways is generated during biosynthesis itself. In the core model, we neglect this ATP produced during biosynthesis by subtracting this from the total amount of ATP required per unit biomass. We then only account for the net ATP requirement ($m_B$) that is provided by one or both of the catabolic modes. Consequently, we regard the respiratory flux in the model as purely catabolic. This results in an underestimation of the predicted specific oxygen uptake flux in comparison to experimental data, since the prediction does not account for oxygen required for ATP production in the TCA cycle during biosynthesis.

To correct for this, we multiply the predicted oxygen uptake flux by $\frac{m^{tot}_B}{m_B}$. This factor represents the total ATP demand ($m^{tot}_B$) over the net ATP demand. This multiplication gives the increased oxygen uptake

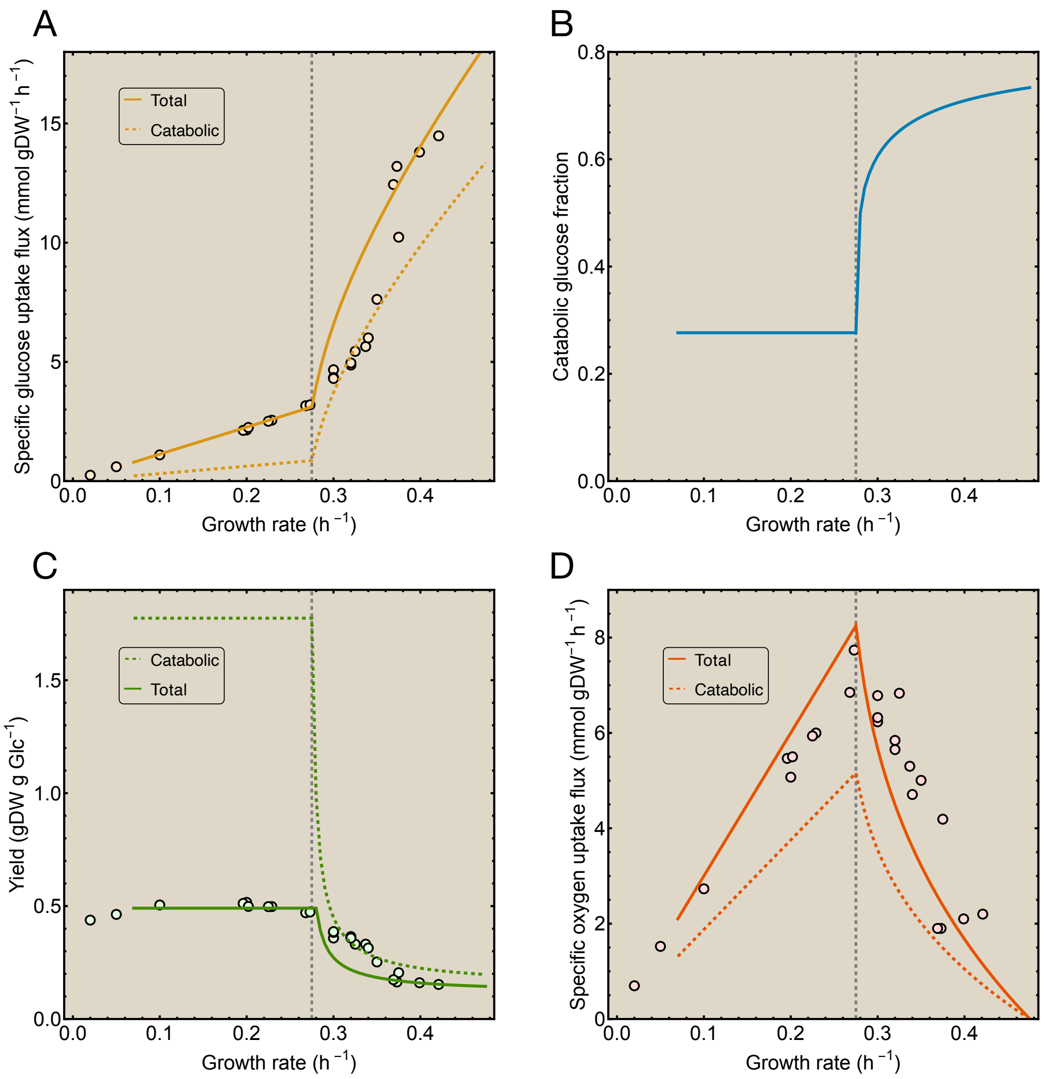


FIGURE S3. **Adjustments for the carbon and oxygen uptake in the yeast model.** A) The total (adjusted) and catabolic specific glucose uptake rate. B) The fraction of glucose consumed by catabolism as function of the growth rate. C) The total (adjusted) and catabolic biomass yield. D) The total (adjusted) and catabolic specific oxygen uptake rate. Chemostat data is reused with permission from [3] (Creative Commons Attribution 4.0 International License).

required to synthesize this extra ATP. This results in the solid red line in Figure S3D, that shows a better correspondence with the experimental data.

### S12.1 Results of the yeast model for total ATP requirement

Another way to improve the prediction for the oxygen uptake is by increasing the net ATP requirement to the total ATP requirement already before solving the optimisation problem in eq. (S30), instead of adjusting this afterwards. Changing this parameter requires a revised calibration of the yeast model. The Mathematica implementation of this model variant is provided in the Supporting Information.

Using the updated calibrated parameters, we again compute the solution to eq. (S30) in terms of the protein fractions. These deviate marginally from the protein fractions found with the net ATP requirement

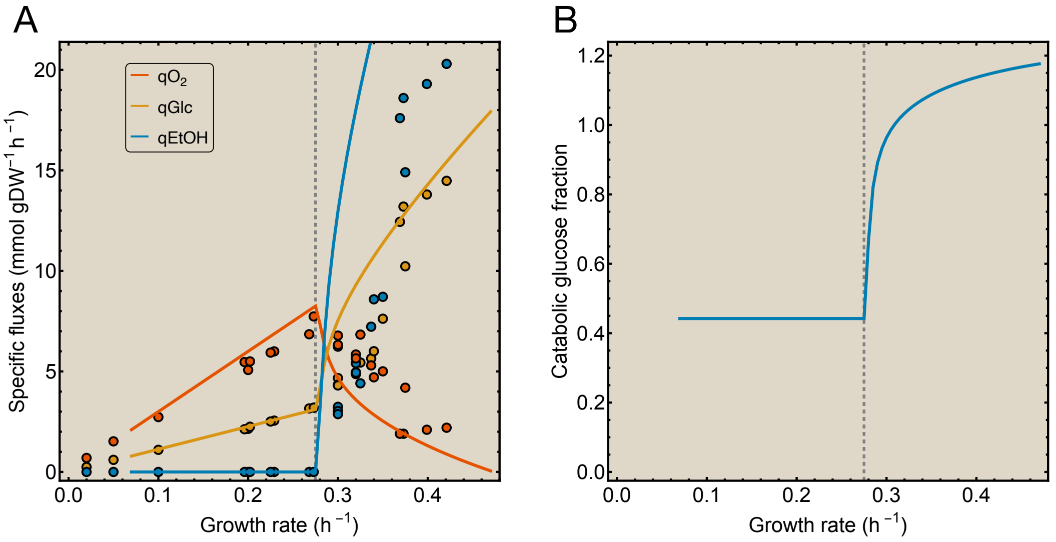


FIGURE S4. **Results of the yeast model accounting for total ATP demand.** A) Specific fluxes predicted by the yeast model together with chemostat data, reused with permission from [3] (Creative Commons Attribution 4.0 International License). B) The fraction of glucose consumed by catabolism as function of the growth rate. This fraction exceeds one when total ATP demand is accounted for.

$m_B$, depicted in Figure 5B. This is because the effect of increasing the ATP demand is (partially) mitigated by the revised calibration.

The results for the specific fluxes are however significantly affected by the increase in ATP demand. A higher ATP demand also requires a higher ATP synthesis flux, i.e., higher glycolytic and respiratory fluxes. An increased respiratory flux results in a higher specific oxygen uptake flux, as represented by the solid red line in Figure S4A. As expected, this also agrees better with the experimental data, similar to the adjustment in Figure S3D.

An increase in the glycolytic flux also requires a higher glucose uptake rate. This partially accounts for the missing carbon required for biomass synthesis, that we corrected for in the previous section. Nevertheless, this does not account for all missing carbon, so an adjustment like in eq. (S43) is still required. We again plot the adjustment factor, given by the catabolic glucose fraction $\frac{S_{cat}}{S_{tot}}$ in Figure S4B. Indeed, this factor is larger compared to Figure S3B for growth rates below the critical growth rate, indicating a smaller adjustment. However, after the critical growth rate the catabolic glucose fraction exceeds one. This would suggest that the catabolic glucose requirement predicted by the model exceeds the measured glucose uptake, which includes both catabolism and anabolism. Such a prediction is unrealistic, but follows directly from the higher ATP demand $m_B^{tot}$. This demand has to be met mostly by ATP generation by glycolysis for growth rates close to $\lambda_{max}$. The low glycolytic ATP yield requires an unrealistically high glycolytic flux. This results in a predicted (catabolic) glucose uptake that exceeds the measured glucose uptake. Because this glycolytic flux equals the fermentation flux (during complete fermentation), this also results in an excessive specific ethanol production flux. This is depicted by the blue line in Figure S4A, which indeed exceeds the measured ethanol production. In reality, part of the ATP synthesis is still performed by respiration during precursor biosynthesis, thereby reducing both the glycolytic and fermentative flux. However, as mentioned before, taking this into account is outside the scope of this model.

To conclude, using the total ATP requirement results in a better prediction for the oxygen uptake flux, but also in unrealistic predictions for other fluxes. Therefore, we argue that analyzing the yeast model with the net ATP requirement and correcting for this afterwards is the better approach. In principle, the core model studies a shift in the catabolic mode and thereby focuses on this part of the metabolic network. This necessarily requires assumptions for the biosynthetic part of the network. These assumptions require adjustments when comparing the model to experimental data.

### S13 Results for the protein fractions in the yeast model

We process the proteomics data from Elsemman et al. [3] to compare with the model results. First, we account for the special case reactions discussed in Section S9. Then, we categorise the reactions according to the pathways and sectors given in Table S1. This is done for all reactions that are assigned to a pathway and for which protein concentrations are measured. Here, we also include measured protein concentrations from batch experiments that were not included to derive the saturation functions in Section S9. We compute the sum of the protein fractions per sector and plot this together with the model predictions as function of the growth rate in Figure 5B.

First, we consider the glycolytic protein fraction. The adjustment to the glucose uptake flux from Section S11 should also be applied to the glycolytic flux, to satisfy the steady-state equations (S25) of the yeast model. Since the mean catalytic constant and saturation of glycolysis are obtained from proteomics data, we can only apply this adjustment by increasing the glycolytic protein fraction. So, we apply the same adjustment to the glycolytic protein fraction in Figure 5B as to the specific glucose uptake flux. Note that the glycolytic protein fraction in Figure 5A is not adjusted, as the protein fractions in the model should add up to one. Despite this adjustment, the glycolytic protein fraction matches the data only qualitatively. This may be due to overestimation of its catalytic constant or saturation function. The glycolytic saturation function vanishes for growth rates below $0.07\,\mathrm{h}^{-1}$, which suggests that it does not accurately represent the mean saturation of glycolysis during slow growth. Because this results in an infeasible optimisation problem, we solve the optimisation problem only for growth rates $\lambda \geq 0.07\,\mathrm{h}^{-1}$. Starting from this value, the glycolytic flux increases with the growth rate. However, the glycolytic protein fraction decreases due to its increasing saturation function, that overcompensates the increased demand in glycolytic flux. Such behaviour is not predicted by the GEM in Elsemman et al. [3], because the authors assume constant saturations.

This behaviour changes after the critical growth rate, because the shift to fermentation requires a higher ATP synthesis rate through glycolysis that can not be compensated by (only) the saturation function. When the growth rate increases further, the glycolytic protein fraction reaches a roughly constant value. This indicates that the increasing ATP demand is completely met by the increasing saturation function in this regime. All these observations are consistent with the experimental data, except for this last effect.

The predicted respiratory protein fraction matches the data quantitatively and shows the expected decrease after the critical growth rate.

For fermentation, we again include only the two canonical fermentation reactions from the proteomics data. Nevertheless, the model prediction does not fit the data well. This may be explained by preparatory expression of the fermentative enzymes and regulation [15; 16]. Similar to glycolysis, also the predicted fermentative fraction stabilises as the growth rate increases. This also follows from its increasing saturation function.

The biosynthetic protein fraction predicted by the model exceeds the experimental data, likely due to an underestimation of the catalytic constant or the (mean) saturation factor. Furthermore, the predicted biosynthetic protein fraction increases hyperbolically with the growth rate, whereas the data seem to show a linear increase. Such a linear relation between the protein fraction and the growth rate indicates a constant saturation factor. However, this does not agree with the saturation function $f_B(\lambda)$ derived from the same proteomics data.

The remaining protein sectors are the $Q$-sector, which is assumed to be constant, and the $NG$-sector. The corresponding protein fractions can not be obtained directly from the proteomics data, since many of these proteins are not assigned to a pathway by Elsemman et al. [3]. Instead, we compute these fractions by subtracting the growth-associated protein concentrations from the total protein concentration. What remains is the growth-unassociated protein, which is the sum of the $Q$- and the $NG$-sector. Therefore, we plot these together in Figure 5B. During slow growth (around $\lambda = 0.07\,\mathrm{h}^{-1}$), the model predicts an increasing growth-unassociated protein fraction, due to the rapidly decreasing glycolytic fraction. For higher growth rates, both the data and the prediction for the growth-unassociated fraction decrease. The model prediction reaches its minimum at the critical growth rate. This minimum corresponds to the (constant) $Q$-sector, which we set to $\phi_Q = 0.25$. In the figure, this is slightly lower due to the adjusted glycolytic protein fraction.

Other factors may also underlie the quantitative discrepancy between the model predictions and the experimental data. Elsemman et al. [3] show that a second metabolic shift occurs in *S. cerevisiae*. Including this results in a better fit of the GEM to the data. This second shift is not contained in our coarse-grained model, as it lacks other proteome sectors that may be constrained. Another effect may arise from the reaction fluxes in terms of averaged kinetic parameters in eq. (31). This approach neglects intrapathway stoichiometry, which also affects the enzyme concentrations.

### S14 Results of the yeast model for constant saturations

Many models for overflow metabolism include phenomenological concentrations and saturations [13; 14], or set saturation factors constant [3; 4; 17]. Here, we account for changing enzyme saturations by including (mean) saturation functions that change with the growth rate. To test if this extra component results in model predictions that agree better with the experimental data, we also compute the solution to the optimisation problem (S30) for constant saturation factors. We compare the results to the same data.

It is not clear *a priori* what is an appropriate constant value for the saturations. Therefore, we implement two model variants with constant saturations. For the first variant, we evaluate the saturation functions at $\lambda = 0.285\,\mathrm{h}^{-1}$, the median of the growth rate of yeast in the chemostat. For the second case, we set all saturations to one, as also done in other models [3; 4]. Both variants of the model are calibrated again using the methods in Section S10, to find the observed values for the critical and maximal growth rates of yeast.

For the first case, a Mathematica implementation is provided in the Supporting Information. Its results are shown in Figure S5. Constant saturations result in protein fractions that vary piecewise linear with the growth rate, as is also illustrated by Figure 2A for the general core model. In general, these predicted protein fractions fit the data less well than the prediction obtained with the saturation functions. The predicted glycolytic protein fraction increases monotonically with the growth rate. This is in better agreement with the data after the critical growth rate, but worse before. The predicted respiratory fraction exceeds the data almost four-fold at some growth rates. The biosynthetic fraction increases linearly with the growth rate, however, without an offset on the y-axis. The predictions for the specific fluxes and the biomass yield fit the data slightly better than the predictions for changing saturations. However, this may be misleading, as for both cases the adjustments from Supplementary Sections S11 and S12 are applied. Since the data for specific fluxes show approximately piecewise linear dynamics, piecewise linear predictions appear to be more accurate after applying these adjustments.

The solution of the second model variant shows behaviour that is very similar to the results in Figure S5. This follows from the model calibration, where we adjust other parameters to find the observed values for the critical and maximal growth rates. This likely compensates for the effect of increasing the saturations from the value of the saturation functions at $\lambda = 0.285\,\mathrm{h}^{-1}$ to one. Since setting saturations constant gives the model less flexibility, this gives similar results for both cases.

Both cases of constant saturations thereby confirm that accounting for changing saturations leads to a better model prediction.

### S15 Analytical solution to the yeast model

The metabolic network of the yeast model in Figure 4 also contains two EFMs, respiration and fermentation. Similar to the general core model analysed in Sections 3.1- 3.2, the steady-state equations (S25) of the yeast model have a unique solution if we consider only one of these EFMs as active mode. We express this solution in terms of the protein fractions and the transporter saturation that change as function of the growth rate.

For respiration, we find

$$
\begin{aligned}
\phi_B^y(\lambda) &= \frac{N_B}{k_{ribo}\langle f(\lambda)\rangle_B}\lambda \equiv C_B^y(\lambda)\lambda \\
\phi_R^y(\lambda) &= \frac{m_B N_R}{(m_R+m_G)\langle k\rangle_R\langle f(\lambda)\rangle_R}\lambda \equiv C_R^y(\lambda)\lambda \\
\phi_G^{R,y}(\lambda) &= \frac{m_B N_G}{(m_R+m_G)\langle k\rangle_G\langle f(\lambda)\rangle_G}\lambda \equiv C_G^{R,y}(\lambda)\lambda \\
f_T^{R,y}(\lambda) &= \frac{m_B}{(m_R+m_G)\langle k\rangle_T\phi_T^y}\lambda.
\end{aligned}
\tag{S44}
$$

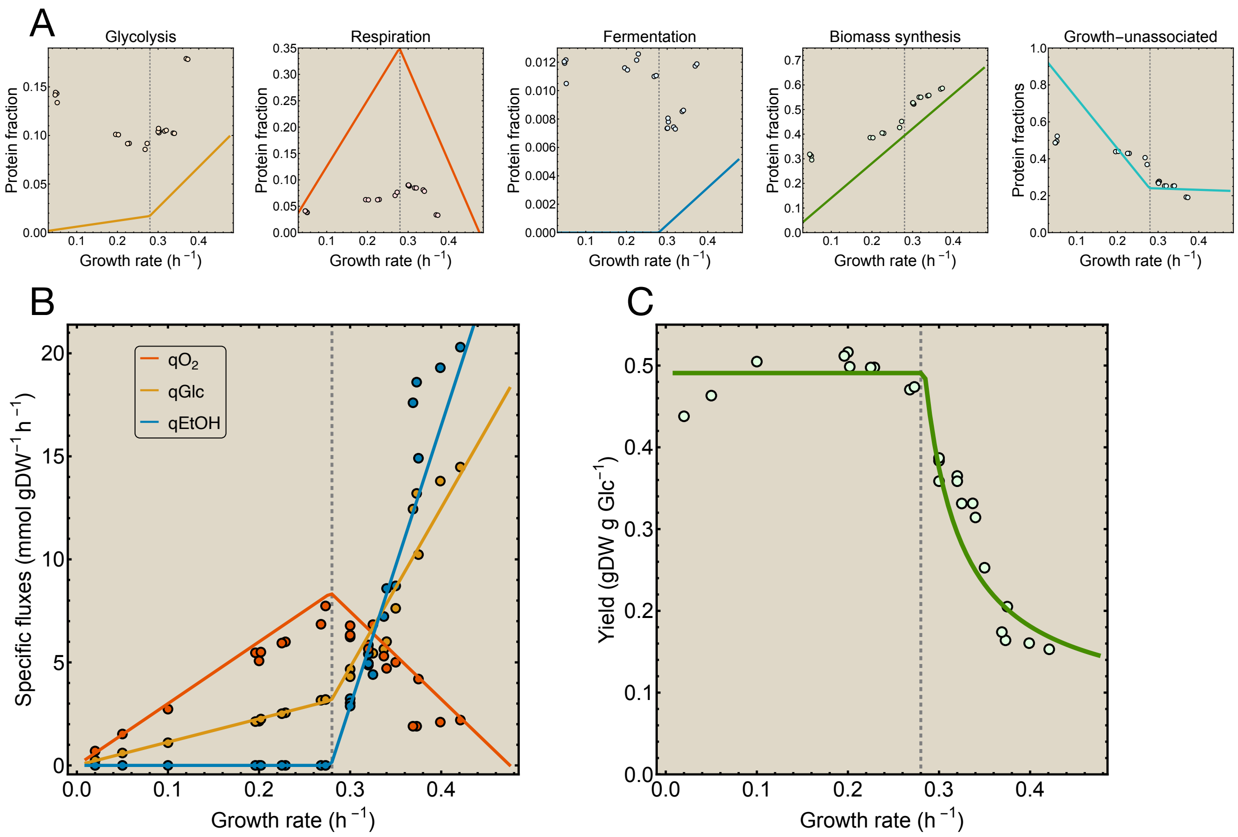


FIGURE S5. **Results of the yeast model for constant saturations factors.** A) Predicted protein fractions for constant saturations together with proteomics data. B) Specific fluxes and C) Biomass yield predicted by the yeast model with constant saturations together with chemostat data. Data is reused with permission from [3] (Creative Commons Attribution 4.0 International License).

Here, $\phi_T^y$ is the constant transporter fraction in the yeast model. The parameter combinations $C_j^y$, that represent the protein costs per process, now change with the growth rate due to the saturation functions. The corresponding growth-unassociated protein fraction

$$\text{(S45)} \quad \phi_{NG}^y(\lambda) = 1 - (\phi_Q^y + \phi_T^y + \phi_B^y(\lambda) + \phi_R^y(\lambda) + \phi_G^{R,y}(\lambda)) = 1 - \phi_Q^y - \phi_T^y - \lambda\left(C_B^y(\lambda) + C_R^y(\lambda) + C_G^{R,y}(\lambda)\right)$$

again decreases with the growth rate, because $e_{tot}$ is assumed fixed. When this protein pool is depleted, yeast reaches its critical growth rate $\lambda_c^y$ and shifts to fermentation. However, obtaining an analytical expression for the critical growth rate by solving $\phi_{NG}^y(\lambda) = 0$ for $\lambda$ is not possible, because the protein costs depend on the growth rate. Instead, we find an approximation for $\lambda_c^y$ by setting the saturation functions constant. Since the observed critical growth rate of yeast during chemostat cultivation is $0.275\,\mathrm{h}^{-1}$, we set the saturation functions constant to their value at $\lambda = 0.275\,\mathrm{h}^{-1}$. This sets the protein costs to $C_j(0.275) \equiv C_j^c$ and gives

$$\text{(S46)} \qquad \lambda_c^y = \frac{1 - \phi_Q^y - \phi_T^y}{C_B^c + C_R^c + C_G^{R,c}}$$

as approximation for the critical growth rate.

For fermentation, the solution to the steady-state equations (S25) is given by

$$\tag{S47} \begin{aligned} \phi_B^y(\lambda) &= \frac{N_B}{k_{ribo}\langle f(\lambda)\rangle_B}\lambda \equiv C_B^y(\lambda)\lambda \\ \phi_F^y(\lambda) &= \frac{m_B N_F}{m_G\langle k\rangle_F\langle f(\lambda)\rangle_F}\lambda \equiv C_F^y(\lambda)\lambda \\ \phi_G^{F,y}(\lambda) &= \frac{m_B N_G}{m_G\langle k\rangle_G\langle f(\lambda)\rangle_G}\lambda \equiv C_G^{F,y}(\lambda)\lambda \\ f_T^{F,y}(\lambda) &= \frac{m_B}{m_G\langle k\rangle_T\phi_T^y}\lambda. \end{aligned}$$

Similarly, we obtain an approximate expression for the maximal growth rate $\lambda_{max}^y$ by evaluating the saturation functions at the observed maximal growth rate, which is $0.47\,\mathrm{h}^{-1}$ for the strain examined in [3]. This sets the protein costs to $C_j(0.47) \equiv C_j^{max}$ and gives

$$\tag{S48} \lambda_{max}^y = \frac{1-\phi_Q^y-\phi_T^y}{C_B^{max}+C_F^{max}+C_G^{F,max}}.$$

The optimisation problem (S30) for the yeast model minimises the transporter saturation $f_T$. The analytical solutions for the transporter saturation for both modes show that $f_T^{R,y}(\lambda) < f_T^{F,y}(\lambda)$ for all growth rates, because the ATP yield of respiration $(m_R+m_G)$ exceeds that of fermentation $(m_G)$. This confirms respiration as the mode that minimises the glucose uptake rate. Evaluating the transporter saturation of respiration at the critical growth rate and that of fermentation at the maximal growth rate yields

$$\tag{S49} \begin{aligned} f_T^c &\equiv f_T^{R,y}(\lambda_c^y) = \frac{m_B}{(m_R+m_G)\phi_T^y\langle k\rangle_T}\lambda_c^y \\ f_T^{max} &\equiv f_T^{F,y}(\lambda_{max}^y) = \frac{m_B}{m_G\phi_T^y\langle k\rangle_T}\lambda_{max}^y. \end{aligned}$$

We combine these equations by computing the ratio of the maximal over the critical transporter saturation, giving

$$\tag{S50} \frac{f_T^{max}}{f_T^c} = \frac{m_R+m_G}{m_G}\frac{\lambda_{max}^y}{\lambda_c^y}.$$

Using the parameters of the yeast model (Table S6), we compute the right-hand side of this equation explicitly, which equals 15.4. In reality, the transporter saturation does not deviate that substantially between the critical and the maximal growth rate during chemostat cultivation [3]. However, the coarse-grained structure of our yeast model directly couples these parameters via eq. (S50). This results in values for the transporter saturation that are unrealistically low (around 0.02) at low growth rates.

### S16 Proteome efficiency in the yeast model

We compute the proteome efficiency of respiration and fermentation in yeast by dividing the ATP synthesis rate by the total protein concentration expended to sustain this ATP flux, similar to the definition in Section 3.3. Using the solutions for the protein fractions for both modes (eq. (S44) and (S47)), this gives

$$\tag{S51} \begin{aligned} \epsilon_R^y(\lambda) &= \frac{m_G v_G(\lambda)+m_R v_R(\lambda)}{e_B^y(\lambda)+e_R^y(\lambda)+e_G^{R,y}(\lambda)} = \frac{m_B}{C_B^y(\lambda)+C_R^y(\lambda)+C_G^{R,y}(\lambda)} \\ \epsilon_F^y(\lambda) &= \frac{m_G v_G(\lambda)}{e_B^y(\lambda)+e_F^y(\lambda)+e_G^{F,y}(\lambda)} = \frac{m_B}{C_B^y(\lambda)+C_F^y(\lambda)+C_G^{F,y}(\lambda)}. \end{aligned}$$

The protein costs related to carbon transport do not appear in the denominator, because the transporter concentration is assumed fixed. Furthermore, the proteome efficiencies in the yeast model depend on the growth rate through the saturation functions. Since these increase monotonically with the growth rate, as demonstrated in Figure S2, also the proteome efficiencies increase monotonically with the growth rate. This is depicted in Figure S6. Here, the efficiency of fermentation abruptly drops to zero at $\lambda = 0.24\ \mathrm{h}^{-1}$, because the saturation of fermentation $(f_F(\lambda))$ vanishes at this growth rate. We consider this as a model artifact.

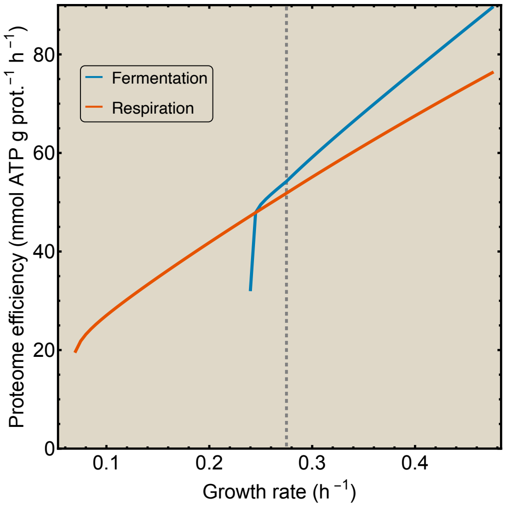


Figure S6. **Proteome efficiencies of both metabolic modes in yeast.** Proteome efficiencies for respiration and fermentation computed as the ATP synthesis rate per total protein concentration expended to sustain the ATP flux. They change as function of the growth rate due to changing saturations. The dashed line represents the critical growth rate.

The figure shows that the proteome efficiency of fermentation exceeds the proteome efficiency of respiration already before the critical growth rate.

Beyond the critical growth rate, respiration can not be used in isolation because of the extra active constraint. So, to compute the respiratory efficiency also in the $\lambda_c \leq \lambda \leq \lambda_{max}$ regime, we had to neglect the constraint that fixes the total protein concentration $e_{tot}$. Therefore, we argue that it is better to evaluate and compare the proteome efficiencies only at the critical growth rate $\lambda_c^y = 0.275\,\mathrm{h}^{-1}$. The corresponding values are given in the first line of Table S5.

As discussed in Supplementary Section S4, a different definition of the proteome efficiency is given by the ATP synthesis rate per amount of catabolic protein expended. For the yeast model, this gives

$$\epsilon_R^{y,cat}(\lambda) = \frac{m_G v_G(\lambda) + m_R v_R(\lambda)}{e_R^y(\lambda) + e_G^{R,y}(\lambda)} = \frac{m_B}{C_R^y(\lambda) + C_G^{R,y}(\lambda)}$$

$$\epsilon_F^{y,cat}(\lambda) = \frac{m_G v_G(\lambda)}{e_F^y(\lambda) + e_G^{F,y}(\lambda)} = \frac{m_B}{C_F^y(\lambda) + C_G^{F,y}(\lambda)}.$$

Evaluating these at the critical growth rate gives the values in the second line of Table S5. This shows that neglecting the biosynthetic protein costs results in much higher values for the proteome efficiency.

These definitions arise from the different interpretations of the protein costs associated with ATP synthesis. For the yeast model, we compute two other values for the proteome efficiency by using the other parameterisation of the ATP requirement for biosynthesis. This is represented by the parameter $m_B$. In the first two lines of Table S5, we computed the proteome efficiencies using the net ATP requirement. The values of the efficiencies using the total ATP requirement for yeast are given in the third and fourth lines. These are computed from the model variant discussed in Supplementary Section S12.

Overall, this results in four different values for the proteome efficiency of both modes in the yeast model. Despite their significant variation, they all meet the condition that fermentation is more proteome-efficient than respiration. Therefore, all these definitions are interchangeable. What matters here are the principles: a shift from respiration towards fermentation due to a limited total protein concentration implies a higher proteome efficiency of fermentation.

The literature values for the proteome efficiency also differ significantly, even when obtained for the same organism. This holds for both *E. coli* [4; 17; 18] and *S. cerevisiae* [3; 6; 17; 18]. In the case of Shen et al. [6], the authors even estimated a higher proteome efficiency for respiration than for fermentation, which violates the condition (S9). Indeed, if overflow metabolism is caused by a limitation on the total proteome, this can not be true. We conjecture that their finding results from an alternative estimate of the protein costs associated with respiration and fermentation. For instance, if protein costs for synthesis of mitochondria are neglected, this results in an increased proteome efficiency for respiration.

| **Definition of proteome efficiency** | **Proteome efficiency respiration** (mmol ATP gram protein$^{-1}$ h$^{-1}$) | **Proteome efficiency fermentation** (mmol ATP gram protein$^{-1}$ h$^{-1}$) |
|---|---|---|
| Net ATP synthesis rate per total growth-associated protein | 51.8 | 54.2 |
| Net ATP synthesis rate per catabolic protein | 346 | 492 |
| Total ATP synthesis rate per total growth-associated protein | 83 | 84.4 |
| Total ATP synthesis rate per catabolic protein | 483 | 533 |

Table S5. **Table with different definitions of the proteome efficiency.** For each definition, the proteome efficiencies for respiration and fermentation in yeast are computed at the critical growth rate $\lambda_c^y = 0.275\,\text{h}^{-1}$. For all cases, fermentation has a higher proteome efficiency than respiration. The values for the proteome efficiencies are computed using the Mathematica implementations of the different yeast model variants, provided in the Supporting Information.

### S17 Break-even analysis for the yeast model

Here, we apply the break-even analysis in terms of parameter perturbations of Section 3.5 and Section S5 to the yeast model. To do this, we should compute $\lambda_{ratio}$, the ratio of the maximal growth rate (eq. (S48)) over the critical growth rate (eq. (S46)). However, because these growth rates are computed at different saturation values, this is no fair comparison of the protein costs. Therefore, we evaluate the saturation functions at $\lambda = 0.37$ hr$^{-1}$, the mean of the critical and the maximal growth rate. The break-even condition then becomes

$$\lambda_{ratio} = \frac{C_B(0.37) + C_R(0.37) + C_G^R(0.37)}{C_B(0.37) + C_F(0.37) + C_G^F(0.37)} = 1. \tag{S52}$$

We reformulate this condition in terms of the model parameters as

$$\frac{N_R}{(m_R + m_G)\langle k\rangle_R \langle f(0.37)\rangle_R} = \frac{N_F}{m_G \langle k\rangle_F \langle f(0.37)\rangle_F} + \frac{m_R N_G}{m_G(m_R+m_G)\langle k\rangle_G \langle f(0.37)\rangle_G}. \tag{S53}$$

We study this condition for five parameter pairs in the yeast model. These correspond to the parameter pairs from the core model that are used to obtain the phase diagrams in Figure 3. For the yeast model, we use its Mathematica implementation given in the Supporting Information, which results in the phase diagrams shown in Figure S7. Figures S7A, B and E depict the results for parameters that appear in both the general and the yeast model. These agree with the results for the general core model, respectively in Figures 3B, C and F.

Figures S7C and D contain the results for perturbation of glycolytic parameters ($k_G$ and $m_G$). Since glycolysis is used by both modes, we expect it to affect both respiration and fermentation. Therefore, we expect results different from those obtained for perturbation of the corresponding fermentative parameters ($k_F$ and $m_F$) in the general core model (see Figure 3D and E). However, also in these cases, the results of both models show qualitative agreement. This indicates that perturbing glycolysis mainly affects fermentation.

An advantage over the phase diagrams in Figure 3 is that we can add a coordinate in Figure S7 that represents the (calibrated) parameter configuration in the yeast model. The distance of this coordinate with respect to the curve (representing the break-even condition (S53)) gives an estimation for the perturbation required to remove overflow metabolism in yeast strains. Parameter pairs for which this distance is (relatively) small may be useful biotechnological targets to manipulate the metabolic shift.

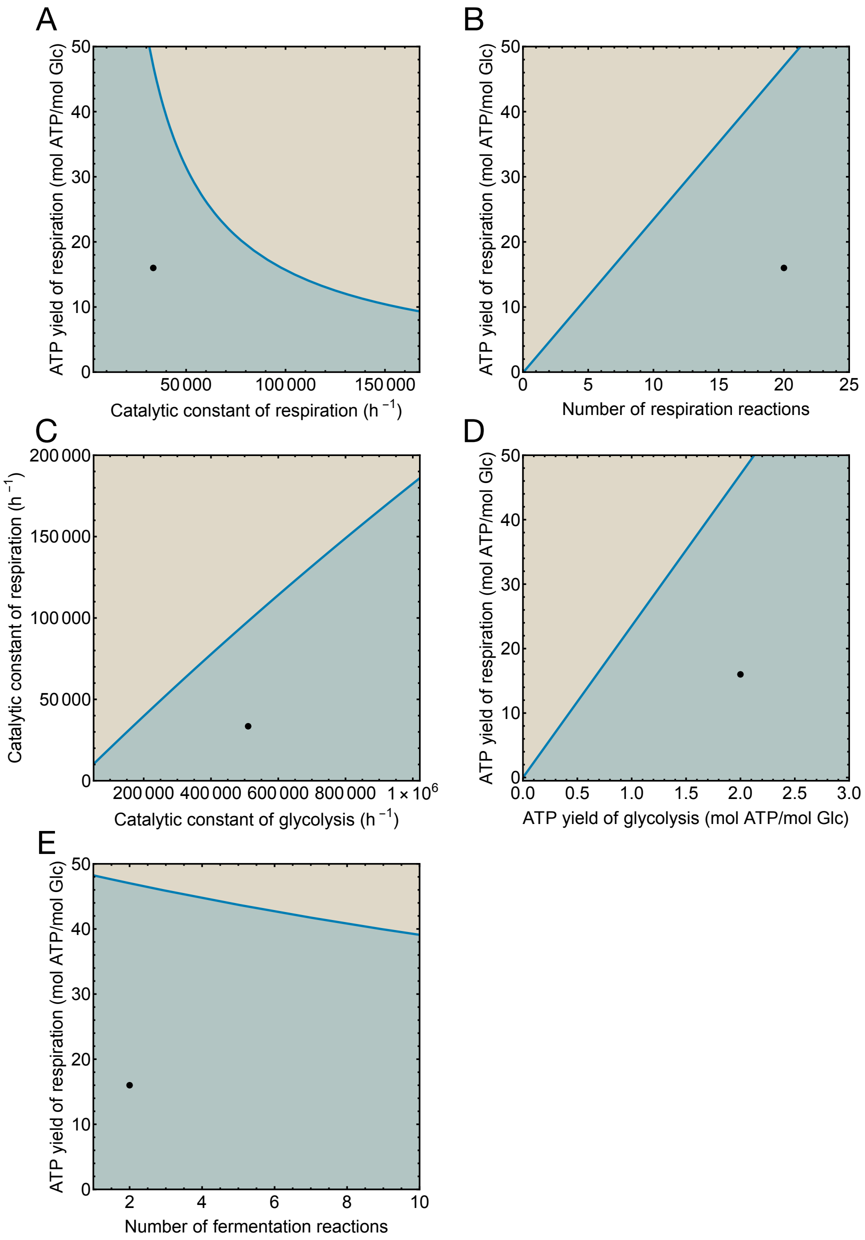


FIGURE S7. **Phase diagrams for the yeast model.** A)-E) The break-even analysis applied to the yeast model in terms of several model parameter pairs. For parameter configurations in the blue region, a metabolic shift occurs. In the brown region a shift will never occur. The black dot represents the parameter configuration used in the yeast model.

| Parameter symbol | Parameter description | Value (source) |
|---|---|---|
| $MW_e$ | Average molecular weight of protein | 50 g/mmol protein ([19]) |
| $N_e$ | Average number of proteins | $2.5 \cdot 10^6$ proteins/$\mu$m$^3$ ([19]) |
| $e_{tot}$ | Average total protein concentration | 4.15 mmol protein/L cells (convert $N_e$) |
| $\phi_T^y$ | Transporter fraction of the total proteome | 0.3% (estimated from proteomics data [3]) |
| $\phi_Q^y$ | $Q$-sector fraction of the total proteome | 25% (estimated from proteomics data [3]) |
| $\rho_e$ | Protein density | $MW_e e_{tot}$ = 208 g/L |
| $\eta_{e/DW}$ | Protein mass fraction per gram dry weight | 40 % ([19]) |
| $\rho_{DW}$ | Dry weight density | $\frac{\rho_e}{\eta_{e/DW}}$ = 519 gDW/L |
| $Y_{ATP/gDW}$ | Net ATP requirement per gram dry weight | 56.3 mmol ATP/gDW ([20]) |
| $m_B$ | Net ATP requirement per mmol protein | $\frac{Y_{ATP/gDW}\rho_{DW}}{e_{tot}}$ = 7038 mmol ATP/mmol protein |
| $Y_{ATP/gDW}^{tot}$ | Total ATP requirement per gram dry weight | 90 mmol ATP/gDW ([20]) |
| $m_B^{tot}$ | Total ATP requirement per mmol protein | $\frac{Y_{ATP/gDW}^{tot}\rho_{DW}}{e_{tot}}$ = 11250 mmol ATP/mmol protein |
| $m_G$ | ATP yield of glycolysis | 2 mmol ATP/mmol glucose |
| $m_R$ | ATP yield of respiration (Oxidative phosphorylation and TCA cycle) | 16 mmol ATP/mmol glucose |
| $Y_{O_2/Glc}$ | Oxygen demand per glucose during respiration | 6 mol $O_2$/mol glucose |
| $Y_{Eth/Glc}$ | Ethanol yield per glucose during fermentation | 2 mol ethanol/mol glucose |
| $k_{ribo}$ | Catalytic constant of the ribosome | 126 h$^{-1}$ ([3]) |

TABLE S6. **Parameters used in the yeast model.** All parameter values are obtained from the literature or computed using literature values.

[6] Yihui Shen, Hoang V. Dinh, Edward R. Cruz, Zihong Chen, Caroline R. Bartman, Tianxia Xiao, Catherine M. Call, Rolf-Peter Ryseck, Jimmy Pratas, Daniel Weilandt, Heide Baron, Arjuna Subramanian, Zia Fatma, Zong-Yen Wu, Sudharsan Dwaraknath, John I. Hendry, Vinh G. Tran, Lifeng Yang, Yasuo Yoshikuni, Huimin Zhao, Costas D. Maranas, Martin Wühr, and Joshua D. Rabinowitz. Mitochondrial ATP generation is more proteome efficient than glycolysis. *Nature Chemical Biology*, March 2024. ISSN 1552-4450, 1552-4469. doi: 10.1038/s41589-024-01571-y. URL https://www.nature.com/articles/s41589-024-01571-y.
